# NUCLEOPORIN1 mediates proteasome-based degradation of ABI5 to regulate *Arabidopsis* seed germination

**DOI:** 10.1101/2023.08.10.552853

**Authors:** Raj K Thapa, Gang Tian, Qing Shi Mimmie Lu, Yaoguang Yu, Jie Shu, Chen Chen, Jingpu Song, Xin Xie, Binghui Shan, Vi Nguyen, Chenlong Li, Shaomin Bian, Jun Liu, Susanne E Kohalmi, Yuhai Cui

## Abstract

NUCLEOPORIN1 (NUP1), a member of the Nuclear Pore Complex (NPC), is located on the inner side of the nuclear membrane. It is highly expressed in seeds; however, its role in seeds including germination has not been explored yet. Here, we identified an abscisic acid (ABA) hypersensitive phenotype of *nup1* during germination. ABA treatment drastically changes the expression pattern of thousands of genes in *nup1*, including the major transcription factors (TFs) involved in germination, *ABI3*, *ABI4*, and *ABI5*. Double mutant analysis of *NUP1* and these ABA-related genes showed that mutations in *ABI5* can rescue the phenotype of *nup1*, suggesting that *NUP1* acts upstream of *ABI5* to regulate seed germination. ABI5, a key negative regulator of germination, is abundant in dry seeds and rapidly degrades during germination. However, its spatiotemporal regulation and interaction with other molecular players during degradation remained to be fully elucidated. We found that NUP1 is physically associated with ABI5 and the 26S proteasome. Mutation in *NUP1* delayed ABI5 degradation through its post-translational retention in nucleolus under abiotic stress. Taken together, our findings suggest that NUP1 anchors the proteasome to NPC and modulates seed germination through proteasome-mediated degradation of ABI5 in the vicinity of NPC in the nucleoplasm.

## Introduction

Seed germination is a critical phase of the plant life cycle initiated by breaking dormancy under favorable conditions. In *Arabidopsis thaliana* (*Arabidopsis*), dry seeds uptake water followed by testa rupture, endosperm rupture, and radicle emergence (Müller et al., 2006; Piskurewicz et al., 2008). Post-germination growth is characterized by cotyledon greening and further root development. All these events are tightly regulated by several internal and external factors such as hormones, light, temperature, and moisture (Finkelstein et al., 2008; Nonogaki., 2017). Optimal germination efficiency under all environmental conditions is a highly desirable trait in agriculture. Therefore, it is important to understand the molecular mechanisms governing seed germination.

Among the factors influencing seed germination, a complex interplay between the phytohormones ABA and gibberellin (GA) has a major impact on the germination process (Liu and Hou, 2018; D. Yang et al., 2022; Zhao et al., 2022). The level of ABA is very high in dry seeds but continuously decreases after imbibition, which is opposite to that of GA. A high level of ABA maintains dormancy while a low level ensures germination (Vishal and Kumar, 2018). In the absence of ABA, PROTEIN PHOSPHATASES TYPE 2C (PP2Cs) inactivate the kinase SNF1-RELATED PROTEIN KINASES (SnRK2s), and downstream genes cannot be activated. When ABA is present, it is detected by the receptor PYRABACTIN RESISTANCE (PYR)/ PYR-LIKE (PYL)/ REGULATORY COMPONENTS OF ABA RECEPTORS (RCAR), then SnRK2s activate several TFs such as ABA Insensitive (ABIs), ABI3, ABI4, and ABI5, for ABA signaling (Park et al., 2009; Cutler et al., 2010; Hauser et al., 2011). ABI5, a member of the basic leucine zipper domain (bZIP) family, is a major TF regulating seed germination (Finkelstein and Lynch, 2000; Kong et al., 2013).

ABI5 is a well-known positive regulator in ABA signaling and plays an important role in seed maturation, germination, and post-germination development (Lopez-Molina et al., 2001). For instance, ABI5 binds to the promoter of genes encoding EMs (Late Embryogenesis Abundant proteins) and affects their expression during the desiccation phase of seed maturation (Carles et al., 2002). ABI5 is a negative regulator of germination; it delays germination when present in excess amount (Skubacz et al., 2016). ABI5 also regulates the expression of *POLYGALACTURONASE INHIBITING PROTEINS* (*PGIPs*) which inhibit the seed coat rupture during germination (Kanai et al., 2010). ABI5 is highly expressed under various abiotic stresses such as osmotic, salt, and drought stresses (Lopez-Molina et al., 2002; Skubacz et al., 2016). Although ABI5 has been extensively studied in recent years (Jiang et al., 2022; Wang et al., 2021; C. Yang et al., 2023), regulation of its temporal and spatial dynamics during the degradation process is poorly understood.

The NPC is one of the largest multi-protein complexes in the cell with a molecular weight of 60-125 MDa (Hampoelz et al., 2019). It has an eightfold symmetrical structure with more than a thousand protein sub-units. It is the only known gateway for the transport of mRNAs and proteins between the nucleus and cytoplasm (Hampoelz et al., 2019; Lin and Hoelz, 2019). *Arabidopsis* NPC contains multiple copies of 30 different types of nucleoporins (Tamura et al., 2010). They are involved in a wide range of molecular and cellular processes, such as *NUP85* and *NUP96* in immune signaling (Gu et al., 2016), and *NUP62* and *SEH1* in auxin response (Boeglin et al., 2016). *Arabidopsis NUP1*, a member of NPC and a putative orthologue of yeast NUP1 was first identified as an interacting protein of the Transcription and Export-2 (TREX-2) complex (Lu et al., 2010). It was later identified as a component of the NPC complex through an interactive mass proteomics approach (Tamura et al., 2010). NUP1 was initially proposed to be a homolog of the animal nucleoporin NUP153; however, there are some differences at the protein domain level (Tamura et al., 2010). In contrast to the NUP153 in vertebrates, *Arabidopsis* NUP1 does not have a DNA-binding domain. NUP153 in mouse embryonic cells regulates stem cell pluripotency through the silencing of developmental genes (Jacinto et a*l*., 2015). *Drosophila* NUP153 regulates gene expression by binding to transcriptionally active regions of the genome (Vaquerizas et al., 2010). However, there is no strong evidence of NUP1 directly regulating gene expression in *Arabidopsis*. We have previously reported the role of NUP1 in mRNA export (Lu et al., 2010) and cell division and expansion (Thapa et al., 2022). Others have reported that *NUP1* is required for maintaining nuclear shape and size (Tamura et al., 2011).

Here, we report that T-DNA insertion lines *nup1-1 and nup1-3* (Supplemental Fig. S1), are sensitive to abiotic stresses during germination. Transcriptomic analysis of *nup1* germinating seeds/young seedlings and genetic analysis showed that *NUP1* is involved in ABA signaling and acts upstream of *ABI5* to regulate seed germination. Further, molecular and cell biology analyses showed that ABI5 is degraded by the 26S proteasome near the vicinity of NPC in the nucleus. Such degradation is delayed in *nup1* plants, leading to the accumulation of ABI5 in the nucleolus and, consequently, delayed seed germination.

## Results

### *NUP1* is required for germination and post-germination establishment

According to public microarray data, *NUP1* is highly expressed in seeds (Supplemental Fig. S2A), and its expression decreases during germination (Supplemental Fig. S2B). This led us to investigate the potential role of *NUP1* in germination. First, the germination efficiency (radicle and cotyledon emergence) of *nup1-1* and *nup1-3* seeds was tested on ½ MS medium agar plates in comparison to the Col-0 wild type. There was no difference in radicle and cotyledon emergence between Col-0 and *nup1-1 or nup1-3* (Fig. 1A-B). However, upon the addition of 1 µM ABA, the cotyledon emergence of *nup1-1 and nup1-3* was delayed by 2-3 days (Fig. 1A, B, D,), but radicle emergence was similar (Fig. 1C). Hereafter, cotyledon emergence was used as a germination indicator in this study. To assess if the mutation in *NUP1* is the real cause behind the ABA-sensitive germination phenotype, we used a previously characterized transgenic line in which the strong *nup1-2* allele was complemented by a *NUP1-YFP* fusion driven by its native promoter *(nup1-2 pNUP1::NUP1-YFP*) (Lu et al., 2010). The delayed germination phenotype of *nup1* was rescued and it’s growth was similar to Col-0 (Fig. 1A-B). This evidence suggests that the ABA sensitive phenotype was indeed due to the mutation in *NUP1*. A *NUP1* overexpression line (*nup1-2 35S::NUP1-YFP*; characterized in Supplemental Fig. S3) was also tested; however, it showed a similar delayed germination phenotype to *nup1-1 and nup1-3* (Fig. 1A-B). The *<u>nup1-</u> <u>1</u>*seed has stronger germination phenotype compared to *nup1-3*, thus we used *nup1-1* for all other downstream experiments. Germination of *nup1-1* seeds was further tested under low exogenous osmotic (sorbitol: 100 and 200 mM) or salt (NaCl: 50 and 100 mM) stresses (Supplemental Fig. S4A-B). While both sorbitol (200 mM) and NaCl (50 and 100 mM) caused a significant reduction in the germination of *nup1-1* seeds, salt stress seemed to have a more detrimental effect (arrested growth) (Supplemental Fig. S4A-D). These data indicate that *NUP1* is required for germination under various abiotic stresses.

**Figure 1.**
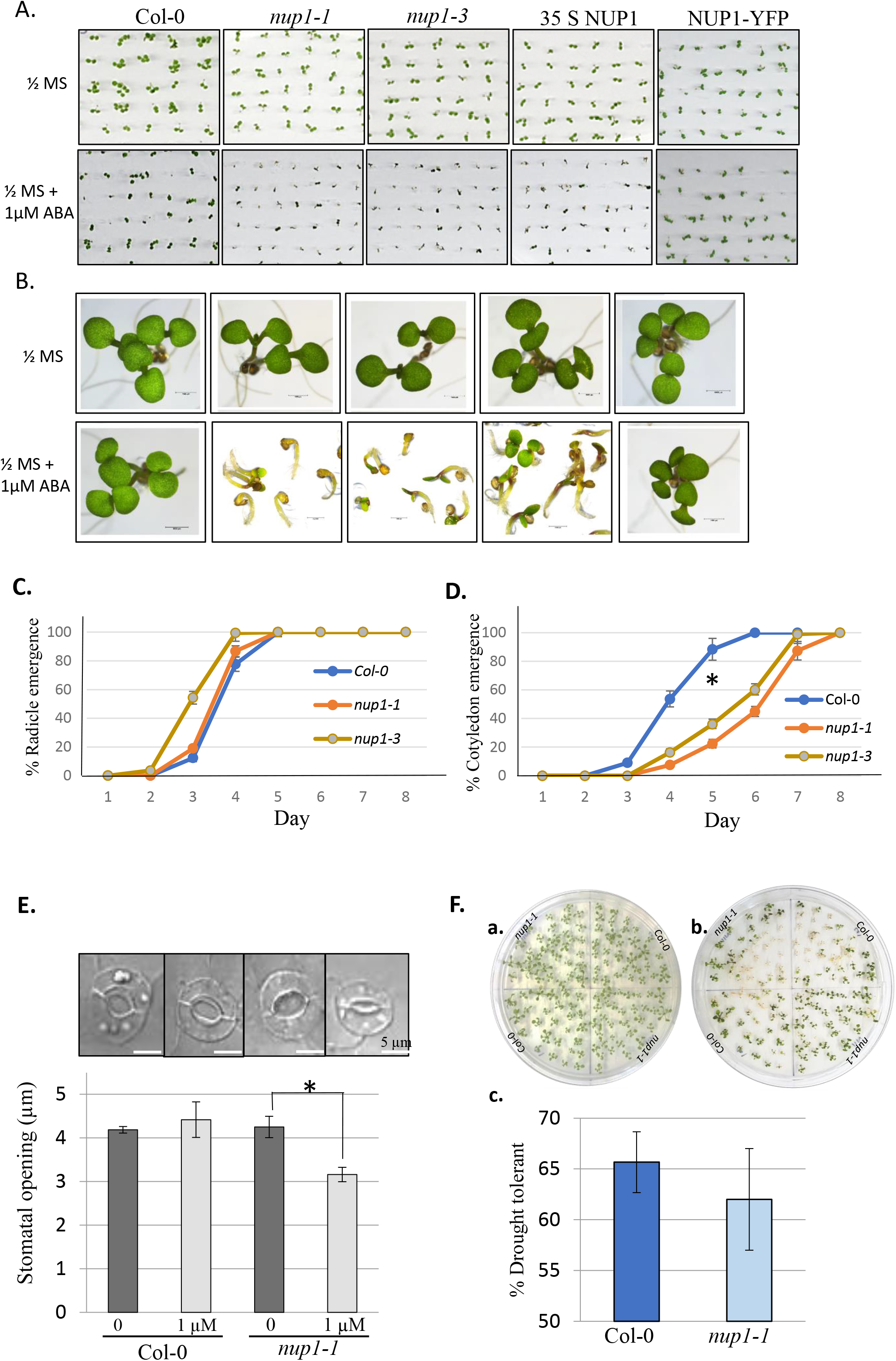
NUP1 is required for germination and early seedling establishment under abiotic stress. Seed germination was recorded for the indicated genotypes grown on ½ MS medium plates with or without 1 µM ABA. Radicles were seen as white tissue, enlarged, and shown in a black box. Values are the mean ± SD. Statistically significant differences were calculated based on Student’s t-tests from day 5 data (* p < 0.05). **A.** Germination of Col-0, *nup1-1, nup1-3,* overexpression line 35S NUP1-YFP (*nup1-2 35S::NUP1-YFP*), and complemented line NUP1-YFP (*nup1-2 pNUP1::NUP1-YFP*) grown on ½ MS medium plates with or without 1 µM ABA. **B.** Representative images of five-day-old seed/seedlings with or without 1 µM ABA **C-D.** Radicle emergence (C) and cotyledon emergence (D) rates were measured from day 1 to 8 of germinating seeds in a growth chamber. **E.** Measurement of stomatal opening in *Arabidopsis* leaves. Leaves of 7-day-old Col-0 and *nup1-1* plants with or without ABA treatment (1 µM) were used to measure the stomatal opening. The stomata opening was significantly reduced in ABA-treated *nup1-1* plants compared to mock-treated *nup1-1*. Corresponding representative stomata samples are shown above the bar graph. Values are mean ± SD from at least 20 samples from three biological replicates. Statistical differences of the stomatal opening between Col-0 and *nup1-1* leaves were determined by Student’s t-test (*p < 0.05). **F**. Drought test assay of *nup1-1* plants under lab conditions. a. Plate at day 7, before exposure to dry air for 6 hrs in the laminar hood. b. Plate after one week of transferring to the growth chamber following drying. c. Comparison of surviving Col-0 and *nup1-1* plants after dry air treatment. No significant difference between the survival rate of Col-0 and *nup1-1* plants. Values are the mean ± SD from at least 30 samples from three biological replicates.

In addition, we measured the stomatal opening in *nup1-1* to understand the effect of *NUP1* mutation on stomatal conductance, a key mechanism mediating plants’ response to various abiotic stresses (Fig. 1E). The stomatal opening of Col-0 and *nup1-1* plants was similar without ABA treatment. In the Col-0 wild-type plants, there was no difference in the stomatal opening with or without ABA treatment; however, in *nup1-1* the stomata were significantly less open after ABA treatment. This result indicates that *nup1-1* seedlings have sensitive stomatal closing. Next, to measure the drought-tolerant capacity, the plants were grown on ½ MS medium plates for 7 days and exposed to airflow in a laminar hood for 6 hrs. Then plants were returned to standard growth conditions for a week, and the percentage of surviving plants was then calculated (Fig. 1F). The mutant plants did not show a drought-tolerant phenotype. Although *nup1-1* seedlings are hypersensitive to stomatal closing, it is not sufficient to make them drought-tolerant. Overall, these data suggest *NUP1* is required for germination and post-germination establishment.

### Differential expression of ABA-related genes in *nup1-1* seeds and seedlings

To investigate the role of *NUP1* in the ABA signaling pathway, the expression level of ABA-related genes in Col-0 wild type and *nup1-1* seeds and seedlings was measured by qRT-PCR. Eight genes from different categories of the ABA core-signaling were selected for the analysis. Seven of them (*ABI3, ABI5, NCED3, NCED5, SnRk2.1, SnRk2.3,* and *SnRk2.6*) were significantly downregulated in *nup1-1* dry seeds compared to Col-0, while *ABI4* expression remained unchanged (Fig. 2A). Similarly, the expression of four genes (*ABI3, NCED3, NCED5*, and *SnRk2.3*) was significantly decreased in 5-day-old *nup1-1* seedlings (Fig. 2B). There was no difference in expression of the other three genes (*ABI5, SnRk2.1*, and *SnRk2.6*) in Col-0 and *nup1-1* 5-day-old seedlings. *ABI4* expression level increased in *nup1-1* 5-day seedlings compared to Col-0. The differences in several ABA-related gene expressions (Col-0 vs *nup1-1*) were higher in seeds compared to seedlings. These results indicate the involvement of NUP1 in the ABA signaling pathway in both seeds and seedlings.

**Figure 2.**
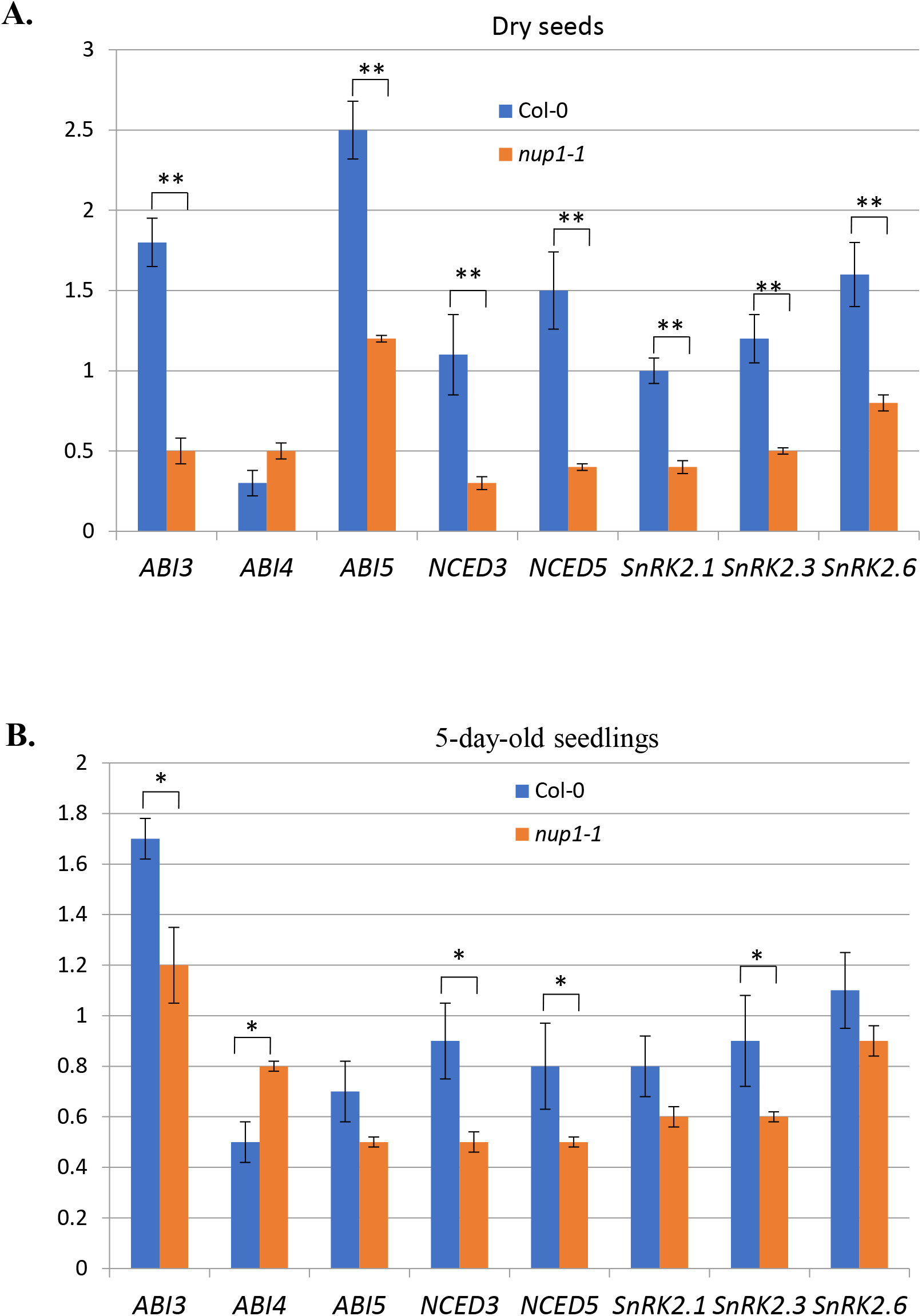
The expression level of ABA-related genes in *Arabidopsis* seeds and 5-day-old seedlings of Col-0 and *nup1-1*. A. Expression of ABA-related genes in Col-0 and *nup1-1* dry seeds. **B.** Expression of ABA-related genes in Col-0 and *nup1-1* 5-day-old seedlings. Gene expression values were normalized with *ACTIN2*. Values are mean ± SEM of three biological replicates and three technical replicates. A two-tailed Student’s t-test was employed to test the statistical difference between Col-0 and *nup1*-*1*. The significant difference is denoted by * for p < 0.05, ** for p < 0.01.

### Transcriptomic analysis of *nup1-1* seedlings reveals changes in the expression of hundreds of genes

Although 5-day-old Col-0 and *nup1-1* seedlings look similar, ABA treatment makes it different (Fig. 1A). At 5 days after sowing seeds on ½ MS medium with ABA, most of the Col-0 seeds were already germinated, and most of the *nup1-1* seeds were arrested at the radicle emergence stage (Fig. 1A). Hence, an RNA-Seq analysis at this stage was expected to capture the difference in the transcriptome, which might explain the ABA-sensitive phenotype of the mutant. Therefore, to understand the changes in the global gene expression pattern in *nup1-1* with and without ABA treatment, an RNA-Seq experiment was conducted for four groups of seedlings: 1). Col-0 wild type; 2). *nup1-1*; 3). Col-0 (ABA) (treated with 1 µM ABA); and 4). *nup1-1* (ABA) (treated with 1 µM ABA). Our results show that, in *nup1-1*, 341 genes were transcriptionally up-regulated and 360 genes were down-regulated compared to Col-0 (Fig. 3A; Supplemental Table S1). In the ABA-treated 5-day-old Col-0, 1018 genes were upregulated while 786 were downregulated compared to mock-treated Col-0 (Fig. 3B; Supplemental Table S2). However, in *nup1-1* (ABA) compared to *nup1-1*, about 4,500 genes in total were differentially expressed (1,692 upregulated and 2,794 downregulated) (Fig. 3C; Supplemental Table S3). There were 599 upregulated and 2,182 downregulated genes in *nup1-1* (ABA) compared to Col-0 (ABA) (Fig. 3D; Supplemental Table S4). Overall, only 701 differentially expressed genes (DEGs) were found between Col-0 and *nup1-1* without ABA treatment. However, the number of DEGs increased after the ABA treatment was much higher for *nup1-1* compared to Col-0. The DEGs were more than two and half times in *nup1-1* (ABA) (4,486 DEGs in Fig. 3C) compared to Col-0 (ABA) (1,804 DEGs in Fig. 3B). The numbers of overlapping and unique DEGs in all four conditions were compared pairwise, and each comparison revealed many shared genes (Fig. 3E). To understand the overall gene expression level, Fragments Per Kilobase of transcript per Million mapped reads (FPKM) was calculated. The average gene expression level (FPKM) for Col-0 and *nup1-1* was similar without ABA treatment, but the expression level significantly increased for both after the ABA treatment (Fig. 3F). Altogether, the transcriptomic profile of *nup1-1* differs from Col-0 and this difference widens more after ABA treatment.

**Figure 3.**
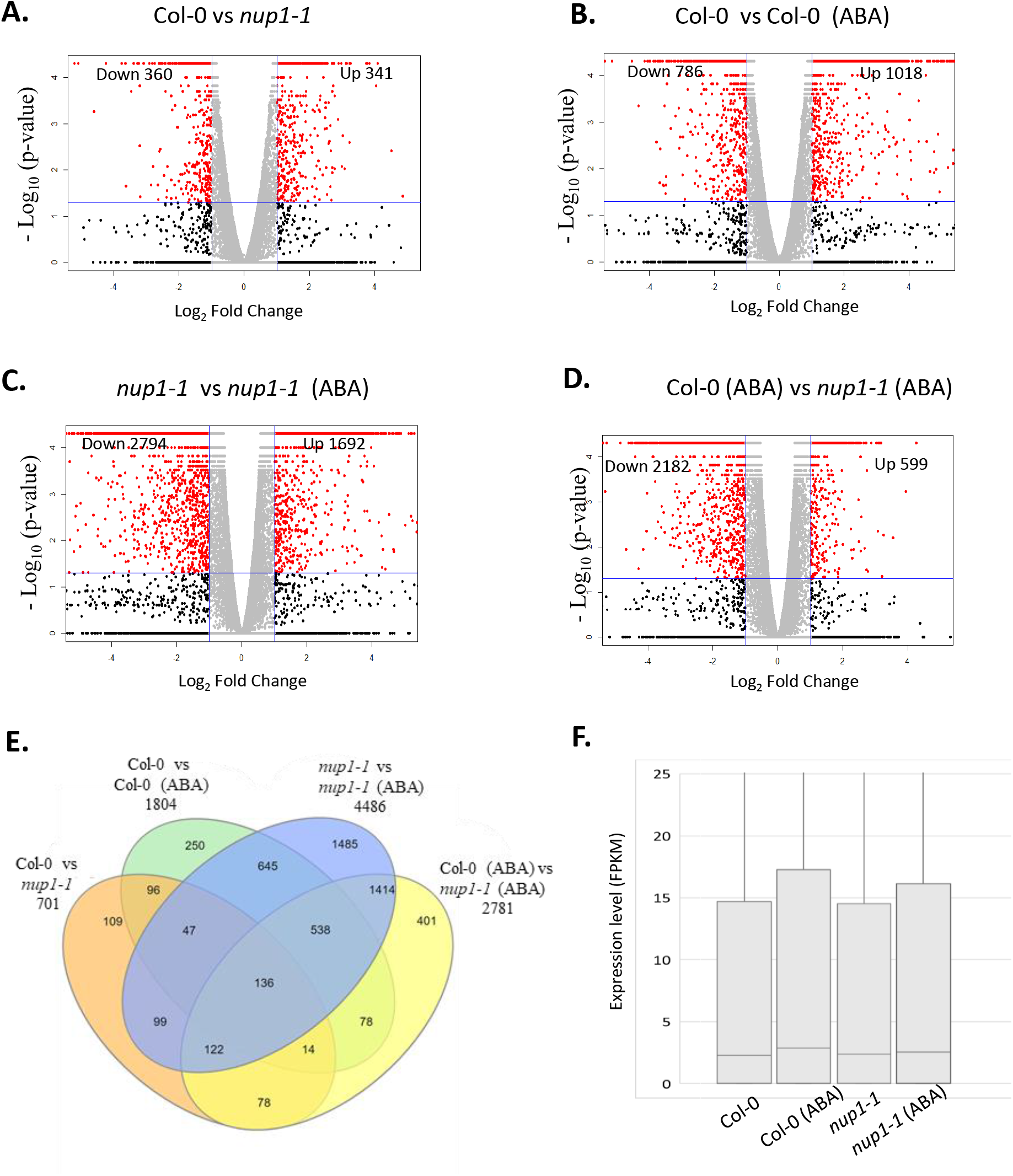
Transcriptomic analysis of *nup1-1* seedlings reveals changes in the expression pattern of hundreds of genes. **A-D.** Volcano plots showing the number of upregulated and downregulated genes in ABA-treated (1 µM) and mock-treated Col-0 and *nup1-1* (specified in each of the panels). X-axis: Log_2_ fold change of Fragments Per Kilobase of transcript per Million mapped reads (FPKM); Y-axis: −Log_10_ (p-value) for the significance of differential gene expression. Red dots represent transcripts that are differentially expressed (p<0.05, fold change >2). Black dots represent transcripts that are not differentially expressed (p ≥0.05, fold change ≥2). Grey represents transcripts that are not differentially expressed (fold change ≤ 2). The horizontal blue line indicates p = 0.05 (upper: p < 0.05; bottom: p > 0.05). The vertical blue lines indicate fold change = 2 (fold change < 2 between two blue lines). **E.** Venn diagram showing the number of overlapped and unique DEGs between different genotypes and ABA treatments. The overlapped and unique number of DEGs (differentially expressed genes) between Col-0 and *nup1-1*, treated with or without 1 µM ABA, were plotted in the Venn diagram. The total DEGs for each comparison are labeled outside the diagram. The total number of overlapped and unique genes is shown inside the diagram. **F.** Box plots showing the average gene expression level. Gene expression level was measured as FPKM for ABA-treated and mock-treated 5-day-old Col-0 and *nup1-1* seedlings. One-way ANOVA was used to determine the statistical difference between different groups. Lowercase letters indicate significant differences between genetic backgrounds. The line inside the box plot represents the median. The interquartile range shows the middle 50% of the data point that ranges between the 25th and 75th percentile. The upper quartile is Q3 and the lower quartile is Q1.

### Gene ontology and differential gene expression analysis uncover key stress-responsive transcription factors upregulated in *nup1-1* seedlings

Gene ontology enrichment analysis (GO-term) of upregulated and downregulated genes between all Col-0 and *nup1-1* was conducted, which highlighted various biological functions (Fig. 4A). Genes involved in multiple pathways related to biotic and abiotic stimulus were upregulated in *nup1-1* compared to Col-0, while only a few pathways were downregulated. Many pathways related to seed development and response to various stimuli were upregulated in Col-0 (ABA) compared to Col-0 while only a few genes and pathways were downregulated. Noticeably, such upregulated pathways were not seen in *nup1-1* (ABA)/*nup1-1*. Overall, *nup1-1* (ABA) has only a few pathways upregulated and most of the pathways downregulated when compared to Col-0 (ABA). This downregulation of a large number of genes and pathways might help explain the delayed growth of *nup1-1* (ABA) germinating seedlings.

**Figure 4.**
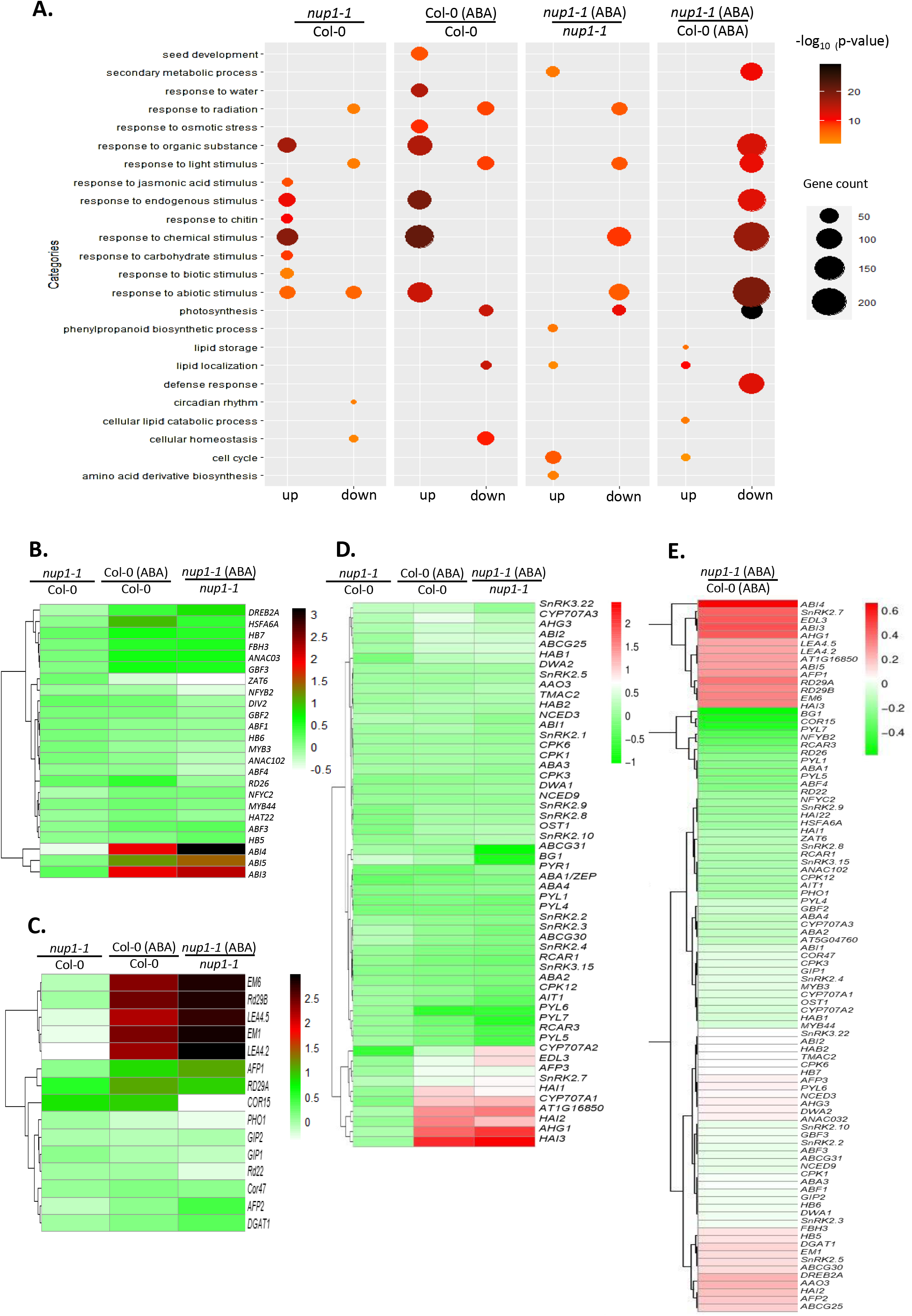
Gene ontology enrichment analysis and Heat maps showing the expression pattern of ABA-related genes in Col-0 and *nup1-1* with or without ABA treatment. **A.** Gene ontology enrichment analysis illustrated as a bubble plot created in R using ggplot2 package. The number of upregulated (up) and downregulated (down) genes for comparison of different groups was plotted as a bubble. The size of the bubble is directly proportional to the number of genes and the color of the bubble represents the −log_10_ p-value as shown in the color key. The heat map was plotted as a −Log_10_ (ratio of FPKM values). Positive and negative values indicate increased and decreased gene expression, respectively. The color key represents the degree of change in gene expression. **B.** Heat map illustrating the expression pattern of selected transcription factors (TFs) in ABA-treated and mock-treated Col-0 and *nup1-1*. Gene expression levels of 24 TFs were plotted for different groups (as labeled). **C.** Heat map illustrating the expression pattern of effector genes in ABA-treated and mock-treated Col-0 and *nup1-1*. Gene expression levels of 15 effector genes were plotted for different groups (as labeled). **D.** Heat map illustrating the expression pattern of ABA-related genes in ABA-treated and mock-treated Col-0 and *nup1-1*. **E.** Heat map illustrating the expression pattern of ABA-related genes in ABA treated *nup1-1* compared to ABA treated Col-0.

To investigate the overall changes in the ABA signaling pathway in mutant seedlings, a comprehensive list of 96 ABA-related genes, divided into nine categories, was compiled from the literature (Supplemental Table S5). One of the important groups is TFs which are normally located and function inside the nucleus, where NUP1 is also located. Therefore, some of them may have a chance of physical interaction with NUP1. Based on the RNA-Seq, expression analysis of 24 TFs shows that most (17/24) of the TFs expression was increased and less than a third (7/24) were decreased in *nup1-1* compared to Col-0 (Fig. 4B). Nevertheless, no single TF expression was particularly highly increased. In Col-0 (ABA) compared with Col-0, the expression of 22 TFs was increased and two were decreased. Of note, the expression levels of three TFs, *ABI3*, *ABI4*, and *ABI5*, were strongly upregulated. In *nup1-1* (ABA) compared with *nup1-1*, expression of 20 TFs was increased and four were decreased. Similar to Col-0 (ABA), the same three TFs were also highly upregulated in this case. Further downstream analysis was, therefore, focused on these three TFs.

To further understand how this three TFs affect the expression of their downstream targets, a list of effector genes (15) co-regulated by *ABI3*, *ABI4*, and *ABI5* were compiled from past studies (Fig. 4C; Supplemental Table S5). In *nup1-1* compared to Col-0, ten genes were upregulated and 5 were downregulated. In ABA-treated Col-0, compared to Col-0, 14 and 1 genes were upregulated and downregulated, respectively. Notably, five genes were highly upregulated (*EM6*, *Rd29B*, *LEA4.5*, *EM1*, and *LEA4.2*; > 15-fold in FPKM and > 1.5 in −Log_10_ values). Similarly, the same five genes were highly upregulated in *nup1-1* (ABA) compared to *nup1-1* (> 30-fold in FPKM and > 2 in Log_10_ values).

Gene expression analysis of all other ABA-related genes (57) was also analyzed in ABA-treated and mock-treated *nup1-1* and Col-0 (Fig. 4D). There was no drastic change in expression in *nup1-1* compared to Col-0. Compared to non-treated, in ABA-treated Col-0 and *nup1-1*, 5 and 7 genes were highly upregulated (> 5-fold FPKM). However, those genes did not fall into any specific functional category or gene family. The changes in expression of all 96 ABA-related genes were analyzed for *nup1-1* (ABA) compared to Col-0 (ABA) (Fig 4E). and it showed that 26 genes were upregulated and 70 were downregulated. Taken together, these analyses indicate that three major TFs (*ABI3*, *ABI4,* and *ABI5*) and their co-target genes were highly expressed in *nup1-1* after ABA treatment.

### Double mutant analysis indicates that *NUP1* acts upstream of *ABI5* in regulating seed germination

From our RNA-Seq data of germinating seedlings, three TFs, i.e., *ABI3, ABI4,* and *ABI5*, are highly upregulated in *nup1-1* compared to Col-0 after ABA treatment (Fig. 4B). We decided to perform a double mutant analysis to delineate the genetic relationship between these genes and *NUP1* in regulating seed germination. To broaden the scope of genetic analysis, two other genes (*ABI1 and ABI2*) that act upstream of the above-selected genes were also included in the genetic analysis. Four double mutants (*nup1-1 abi1-2*, *nup1-1 abi2-2*, *nup1-1 abi3-7,* and *nup1-1 abi5-8*) were generated by crossing the corresponding single mutant plants and subsequent genotyping for T-DNA insertion. Double homozygous *nup1-1 abi4-2* could not be obtained, therefore not included in this study.

Seeds of Col-0, *nup1-1*, the four *abi* single mutants, and the four double mutants were grown on ½ MS medium with or without 1 µM ABA (Fig. 5; Supplemental Fig. S5). Seeds of all genotypes show similar germination and growth on ½ MS (control) (Fig. 5A; Supplemental Fig. S5A, C, E). Also, all the seeds have a similar rate of radical emergence. However, there were clear differences in cotyledon emergence (germination marker in this study) between different genotypes of seeds treated with ABA (Fig. 5B, C; Supplemental Fig. S5B, D, F, G, I, K).

**Figure 5.**
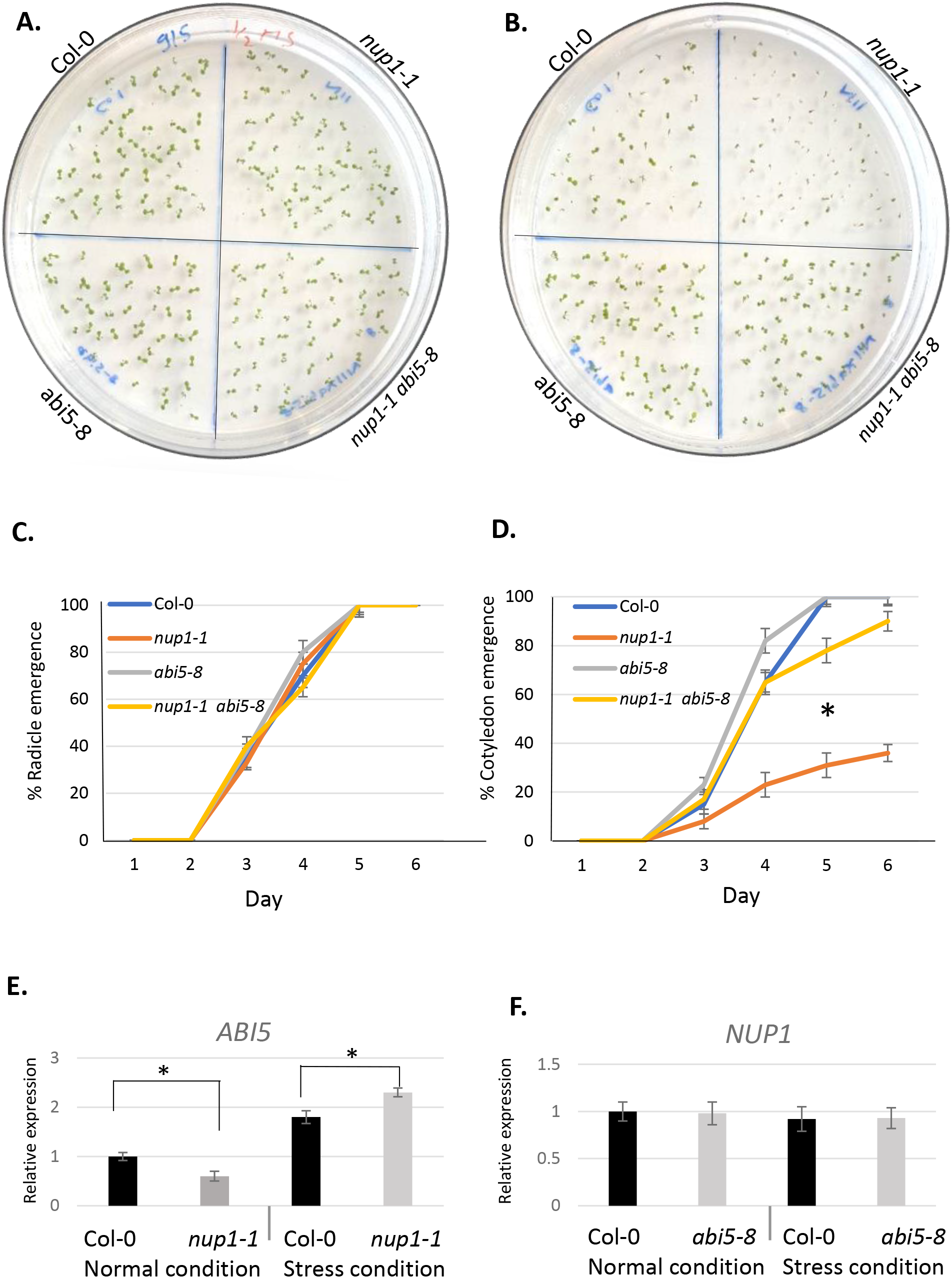
*NUP1* acts upstream of *ABI5* in regulating *Arabidopsis* seed germination. **A-B.** Seed germination, recorded at day 5, of Col-0, *nup1-1*, *abi5-8*, and *nup1-1 abi5-8* sown and grown on ½ MS medium plates as a control (A) and on ½ MS medium plates with 1 µM ABA as a treatment (B). **C-D.** Graph showing the percentage of radicle and cotyledon emergence (germination) of seeds from different genetic backgrounds grown for 5 days on ½ MS medium plates with 1 µM ABA. The *nup1-1* germination is significantly lower than Col-0*, abi5-8,* and *nup1-1 abi5-8* (Student’s t-test, * p < 0.05). The germination of *nup1-1 abi5-8* is similar to Col-0. **E-F.** *ABI5* and *NUP1* expression levels in 5-day-old *nup1-1* and *abi5-8* plants, respectively, with or without stress condition (1 µM ABA for 3 hrs). The gene expression levels were normalized to *ACTIN2*. Values are the mean ± SD of three biological replicates. The statistical differences in germination were calculated from day 5 data using a two-tailed Student’s t-test, * p < 0.05.

Under ABA treatment, *abi1-2* and *abi2-2* seedlings showed delayed germination compared to Col-0 (Supplemental Fig.S5B, D, H, K). However, *abi3-7* and *abi5-8* have a germination rate similar to that of Col-0 even after ABA treatment (Fig. 5B and 5D; Supplemental Fig.S5F, L). Out of the four double mutants, only *nup1-1 abi5-8* could rescue the delayed germination (cotyledon emergence) phenotype of *nup1-1* (Fig. 5B and 5D). All other three double mutants showed ABA sensitive phenotype during germination, which is similar to *nup1-1*. These genetic results suggest that *NUP1* acts upstream of *ABI5* during seed germination in the ABA signaling pathway.

To further support the notion of *NUP1* acting upstream of *ABI5*, *NUP1* expression in *abi5-8* seedlings and *ABI5* expression in *nup1-1* were measured (Fig. 5E-F). Under normal conditions, the *ABI5* expression was initially significantly lower in *nup1-1* compared to Col-0; however, after 1 µM ABA treatment (a stress condition), *ABI5* expression became significantly higher (Fig. 5E). In contrast, the expression level of *NUP1* was relatively constant with or without 1 µM ABA treatment in *abi5-8* seedlings compared to Col-0 (Fig. 5F). These gene expression data also strengthen the claim of *NUP1* being genetically upstream of *ABI5*. This raised the possibility of NUP1 binding to the promoter of ABI5 to regulate its expression. However, a recent study showed that NUP1 does not bind to the promoter of *ABI5* or any other genes for their regulation (Bi et al., 2017). Therefore, we focused our attention on the potential physical interaction between NUP1 and ABI5.

### NUP1 interacts with ABI5 and the proteasome in the nuclear basket region to maintain ABI5 homeostasis post-translationally

To study the interaction between NUP1 and ABI5, two assays (co-localization and BiFC) were conducted. First, for co-localization study, a plasmid expressing an ABI5-CFP fusion under the *35S* promoter (*35S::ABI5-CFP*) was constructed and transiently expressed in leaves of 5-day-old *nup1-2 pNUP1::NUP1-YFP* seedlings. As expected, NUP1 was expressed around the nuclear envelope, while ABI5 was expressed inside the nucleus (Fig.6A and B, respectively). Importantly, as shown in Figure 6C (highlighted by the red box), there is a co-localization (orange color) of these two proteins in the nuclear envelope. Second, the protein interaction was tested through the BiFC assay. The interaction between NUP1 and ABI5 was detected by confocal microscopy (Fig. 6D).

**Figure 6.**
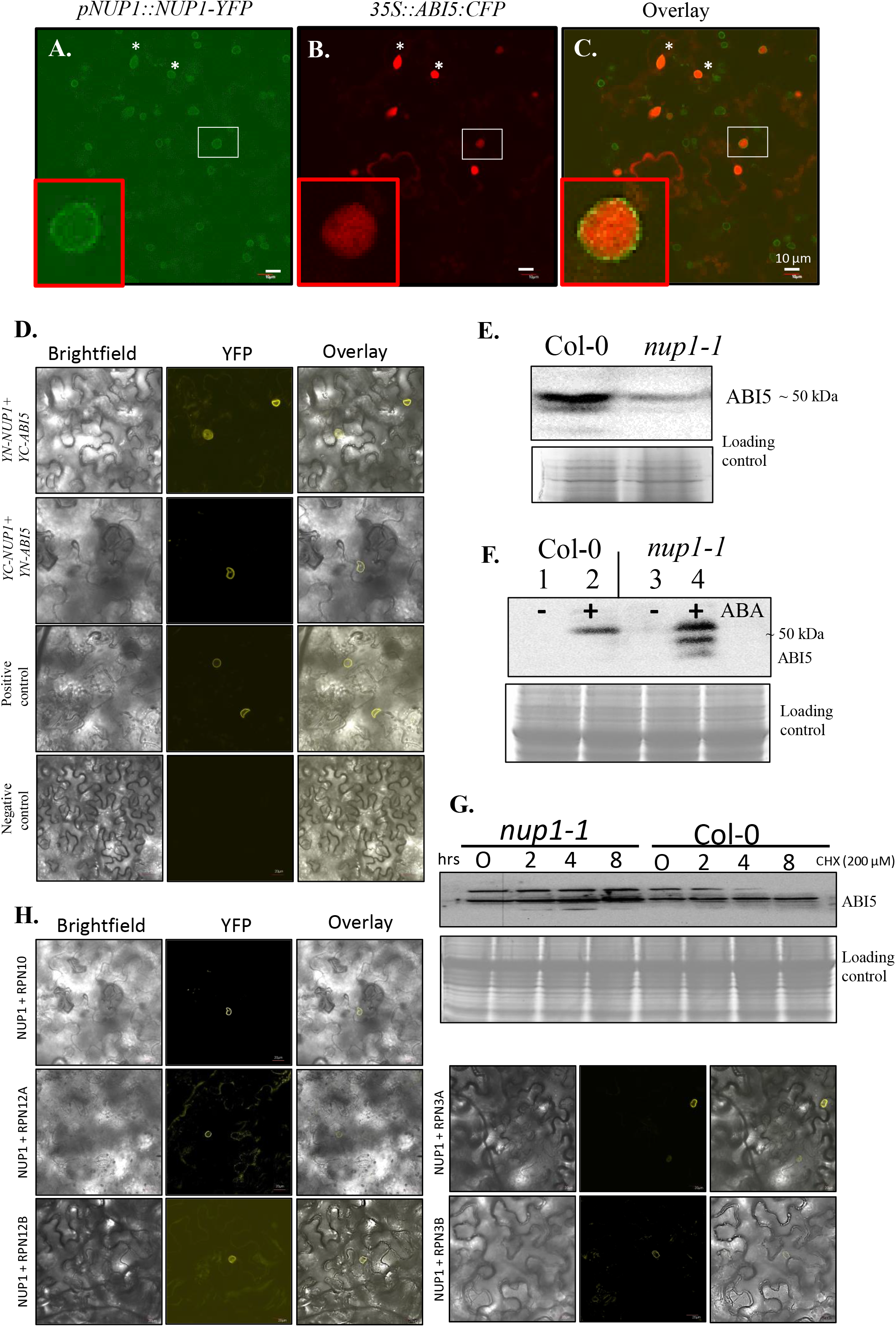
Physical interaction of NUP1 with ABI5 and the proteasome and the effect of NUP1 on the ABI5 level. **A-C**. Co-localization assay of NUP1 and ABI5 in *Arabidopsis*. Four-day-old *Arabidopsis* transgenic seedlings (*nup1-2 pNUP1::NUP1-YFP*) were infiltrated for transiently expressing an ABI5-expressing plasmid (*35S::ABI5-CFP*). After 2 days, leaves were observed under a confocal microscope for co-localization signals. NUP1-YFP signal detected in A; ABI5-CFP signal detected in B, and C is an overlay of A and B. The small red boxes in A-C are enlarged signals of the indicated proteins (white boxes). Co-localization of NUP1-YFP and ABI5-CFP can be seen as orange color in nuclear envelope regions (red box of Figure C). **D.** BiFC assay testing the interaction between NUP1 and ABI5. YFP signal was detected by confocal microscopy in tobacco leaves. *Agrobacterium* containing BiFC vectors (YN-NUP1+YC-ABI5) and (YC-NUP1 + YN-ABI5) were used to transform 4-week-old tobacco leaves. After 2 days, the leaves were observed under a confocal microscope for potential protein interaction signals. Positive control: interaction between NUP1 and THP1 (Lu et al., 2010); negative control:YN-NUP1 and YC-CENTRIN2 (Zhang et al., 2020). **E-F.** Western blot showing the ABI5 levels in dry seeds and 5-day-old seedlings. Detection of ABI5 in protein extracts from Col-0 and *nup1-1* seeds (E) and seedlings (F) (treated with or without 1 μM ABA (+/-ABA) for 5 days). Antibody used: anti-ABI5 (ab98831). ABI5 has three splicing isoforms which may result in up to three bands in Western blot. Scale bar: 10 μm. Coomassie blue-stained gel image was used as a loading control. **G.** Western blot showing the level of ABI5 in CHX (200 μM) treated Col-0 and *nup1-1* seedlings over the indicated time. Coomassie blue-stained gel image was used as a loading control. **H.** BiFC assay testing the interaction between NUP1 and five proteins from the 26S proteasome.

To understand the role of ABI5 in seed germination of *nup1-1*, the abundance of ABI5 in *nup1-1* seeds and seedlings was analyzed by Western blot. The ABI5 level was higher in Col-0 compared to *nup1-1* in dry seeds (Fig. 6E), which is in agreement with the transcript level from qRT-PCR (Fig. 2 A). ABI5 was not detected in 5-day-old Col-0 and *nup1-1* seedling (Fig. 6F, lane 1, 3), which was expected, as ABI5 level decreases rapidly during the seed to seedling transition (Finkelstein and Lynch, 2000). However, ABI5 was detected after ABA treatment in both Col-0 and *nup1-1* seedlings (Fig. 6F, lane 2, 4). The level of ABI5 was higher in *nup1-1* seedlings compared to Col-0. This increased level of ABI5 in mutant seedlings may be due to an increase in transcription of *ABI5* and/or delay in the degradation of ABI5. The possibility of NUP1 directly regulating the *ABI5* gene expression could be ruled out according to a recent genome-wide occupancy study (Bi et al., 2017). Therefore, the prospect of delayed degradation of ABI5 in *nup1-1* seedling was further explored here by a translational inhibitor (Cycloheximide, CHX) treatment assay. The Col-0 and *nup1-1* seedlings grown under ABA stress were transferred to CHX-supplemented agar plates and treated for 2 to 8 hrs. After CHX treatment, the ABI5 protein level decreased rapidly in Col-0 seedlings but remained stable in *nup1-1* (Fig. 6G). The translational inhibitor treatment experiment suggests a post-translational regulatory mechanism by which NUP1controls ABI5 abundance.

Since NUP1 has some role in maintaining ABI5 homeostasis, we looked further into the factors linking these two proteins during degradation. Our earlier studies suggest that NUP1 tethers the 26S proteasome to NPC through the TREX-2 complex (Lu et al., 2010; Tian et al., 2012); and other studies have shown that ABI5 is degraded through the 26S proteasome (Lopez-Molina et al., 2001; Liu and Stone, 2010). Thus, we hypothesized that NUP1 facilitates the 26S proteasome-based degradation of ABI5 in the nuclear basket region. In a recent study, an immunoprecipitation mass spectrometry experiment revealed the physical interaction between NUP1 and 19 members of the 26S proteasome complex (Zhang et al., 2020), confirming our earlier work. To further confirm these results, we performed a BiFC assay between NUP1 and five proteins from the proteasomal complex (RPN12a, RPN12b, RPN3a, RPN10, and RPT2B). All five proteins interacted with NUP1 in nuclear envelope regions (Fig. 6H). Together, these findings support a scenario that NUP1 interacts with ABI5 and the 19S RP of the proteasome in the nuclear basket region to maintain the ABI5 level.

### ABI5 is predominantly localized in nucleoplasm but mutation in *NUP1* leads to its nucleolar retention under abiotic stress

Our protein-protein interaction data suggest the NPC basket region as a potential ABI5 degradation site. Although ABI5 is known to localize inside the nucleus (Liu and Stone, 2013), its subnuclear distribution has not been well-studied. Here, we revisited the issue by using several chemical and biological markers. First, we used the nuclear marker propidium iodide (PI) to determine the region of the nucleus stained by it (Fig. 7A). PI stained nucleolus Fig. (7Aa) and/or cell wall (Fig. 7Ab). To confirm these results, we also used another nuclear marker DAPI (Fig. 7Ac). Together, these visualization studies strongly show that PI stains nucleolus and /or cell walls. We thus decided to use PI as a nuclear marker in the following experiments because it stains both nucleolus and cell wall, which is desirable for the visualization of subcellular compartments. Also, PI is less toxic to cells compared to DAPI which may dampen the ABI5-GFP fluorescence signal in a longer experimental setup.

**Figure 7.**
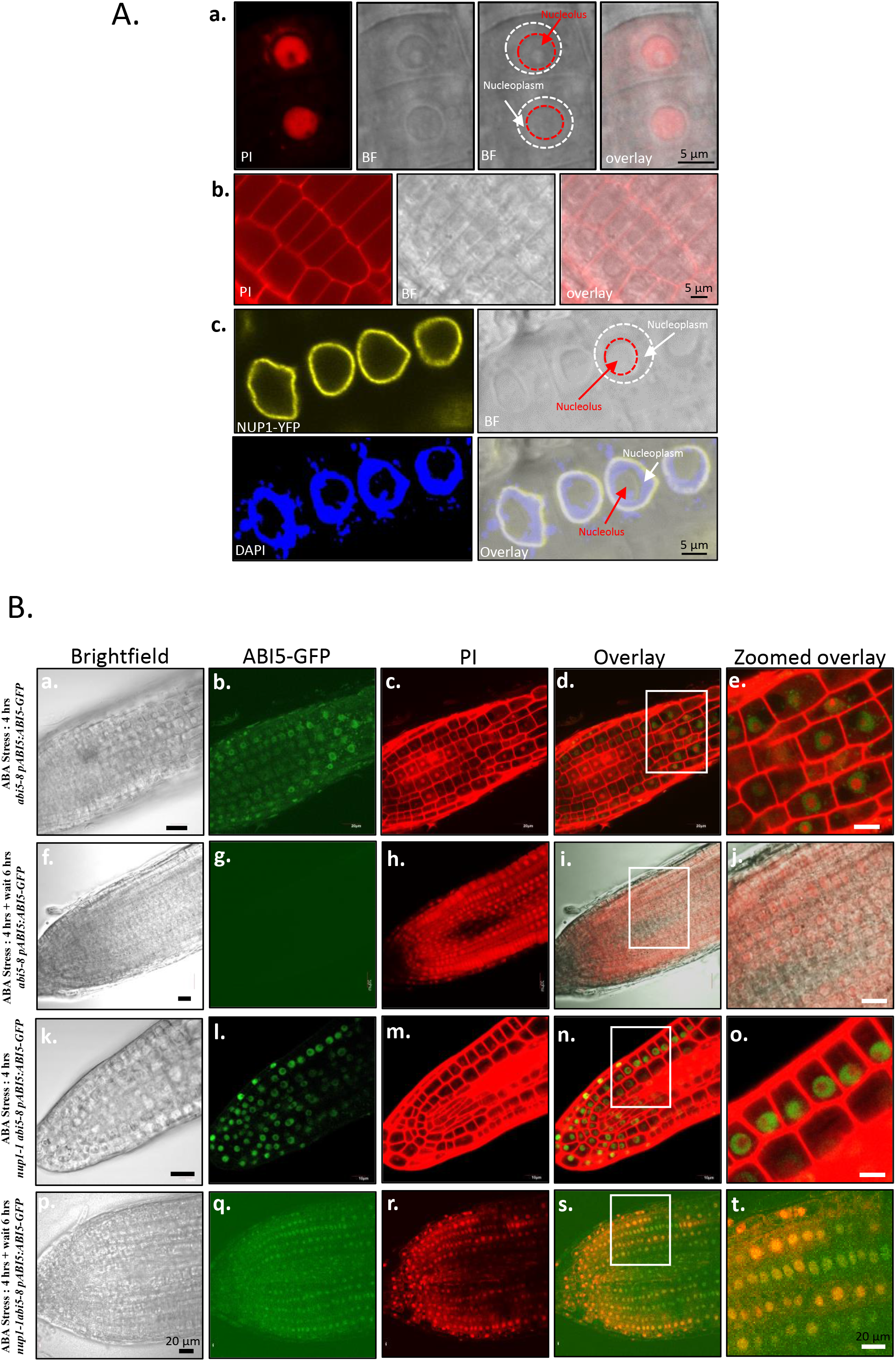
Tracking the subcellular localization of ABI5 expression and degradation in *Arabidopsis* roots. A. Use of propidium iodide (PI) to label the nucleus in *Arabidopsis* root. a. PI labeling of *Arabidopsis* root showing stained nucleus. PI: propidium iodide; BF: Brightfield. Brightfield image is marked to show nucleolus and nucleus boundary. The big white circle represents the nuclear envelope and the red small circle represents the nucleolus boundary. b. PI labeling of *Arabidopsis* root showing labeled cell wall. c. Transgenic *Arabidopsis* expressing *pNUP1::NUP1-YFP*. The NUP1-YFP marker is combined with DAPI to confirm the boundary of nucleus and nucleolus in a plant root cells. B. Tracking subcellular localization of ABI5 expression and degradation by microscopy. a-e. Confocal images showing nuclear (nucleoplasmic) localization of ABI5-GFP signals in *abi5-8 pABI5::ABI5-GFP* plants, treated with 20 µM ABA for 4 hrs and immediately observed. f-j. Confocal images showing absence of ABI5-GFP signals in *abi5-8 pABI5::ABI5-GFP* plants, treated with 20 µM ABA for 4 hrs and observed after another 6 hr wait period. k-o. Confocal images showing nuclear (nucleoplasmic) localization of ABI5-GFP signals in the *nup1-1 abi5-8 pABI5::ABI5-GFP* plants, treated with 20 µM ABA for 4 hrs, and immediately observed. p-t. Confocal images showing nuclear (nucleolus) localization of ABI5-GFP signals in *nup1-1 abi5-8 pABI5::ABI5-GFP* plants, treated with 20 µM ABA for 4 hrs and observed 6 hrs later after removal of ABA. Brightfield: a, f, k, p. GFP channel for ABI5 localization: b, g, l, q. PI channel for nuclear marker: c, h, m, r. Overlay: d, i, n, s; and enlarged version of the sections marked by the white boxes displaying co-localization of GFP signals and PI signals (orange color): e, j, o, t. Scale bar: 20 µm.

To investigate the ABI5 localization at a subnuclear level and the role of NUP1 in the ABI5 degradation pathway, two transgenic lines were generated: one for expressing an ABI5-GFP fusion driven by its native promoter in the *abi5-8* background (*abi5-8 pABI5::ABI5-GFP*), as a control; and the other experimental line expressing the same transgene, but in the *nup1-1 abi5-8* double mutant (*nup1-1 abi5-8 pABI5::ABI5-GFP*). In a preliminary experiment, we determined the time frame of the expression and degradation of ABI5-GFP for visualization using confocal microscopy. It took about 4 hrs of ABA treatment (20 µM) to express enough ABI5 for detection and another 6 hrs for near complete degradation of ABI5 (Supplemental Fig. S6). Based on these results, the control plants were first treated with 20 µM ABA for 4 hrs, and half of them were immediately observed under a confocal microscope (Fig. 7B a-e). The remaining control plants were transferred to a standard medium for another 6 hrs before being observed (Fig. 7B f-j). The immediately observed control plants showed mostly nucleoplasmic localization of ABI5 (Fig. 7B a-e), while those observed 6 hrs later showed no ABI5 expression (Fig. 7B f-j). For the experimental transgenic line in the double mutant background, the same procedures were followed to investigate the role of NUP1 in ABI5 localization. The immediately observed experimental plants showed mostly nucleoplasmic localization (Fig 7B k-o), and those observed 6 hrs later showed nucleolar localization (Fig. 7B p-t). Overall, these results strongly suggest that ABI5 is predominantly localized in the nucleoplasm under abiotic stress and becomes degraded there gradually after the removal of stress. Interestingly, in *nup1-1* plants, ABI5 is retained in the nucleolus instead of being degraded in the nucleoplasmic region.

## Discussion

We investigated the function of *NUP1* during seed germination under various abiotic stresses. The findings that *nup1-1* and *NUP1*-overexpressed seeds show delayed germination suggest the need for a wild-type level of NUP1 for proper germination. The *nup1-1* seedlings were not drought tolerant, the reason for which might be that NUP1 also plays a major role in many other basic cellular activities such as mRNA export, nuclear size regulation, and other transport activities (for review, see Tamura, 2020).

Our data about the proteasome being present in the nucleus and tethered to NPC are consistent with studies in other organisms. Fluorescence correlation spectroscopy analysis revealed that the yeast nucleus has about five times more proteasome than the cytoplasm (Pack et al., 2014). A recent study in *Chlamydomonas reinhardtii* using cryo-electron tomography imaging reported that proteasomes are densely accumulated at two distinct sites in the nucleus, the NPC basket, and the nuclear membrane surrounding NPC (Albert et al., 2017). In *Schizosaccharomyces pombe*, nucleoporin TPR of the nuclear basket is required for localization of proteasomes to the nuclear membrane (Salas-Pino et al., 2017). These studies provide convincing evidence of proteasomes being present around NPC in several organisms. In *Arabidopsis*, a recent study demonstrated the strong interaction between NUP1 and the 26S proteasome (Zhang et al., 2020). Nineteen members of the 19S Regulatory Particle (RP) were detected to be co-immunoprecipitated with NUP1. Interestingly, no proteins of the 20S Core Particle (CP) were identified in the process (Zhang et al., 2020). This suggests that NUP1 may be the linkage connecting proteasome to NPC through the 19S RP (lid and base) of the proteasome. In our study, the physical interaction of NUP1 and ABI5 was detected only by BiFC but not by Co-IP. This might be a consequence of the weak and transient nature of the interaction or the large size of the associated protein complexes including TREX-2 and other adapter proteins, which could lower the efficiency of Co-IP. Interestingly, a recent study using a more sensitive technique demonstrated that NUP1 was pulled down by ABI5-TurboID-GFP in an IP-MS experiment (Log_2_ (FC (ABI5-TurboID-GFP vs TurboID-GFP)) = 2.59 and p-value < 0.05) (Yang et al., 2023). This finding is consistent with our BiFC result (Fig. 6D), and together they strongly support a physical association between NUP1-ABI5, which is likely indirect and transient in nature. Of note, the plant materials used by Yang et al were also young seedings treated with ABA; and NUP1 is the only member of NPC among the 67 proteins pulled down by ABI5 (Yang et al., 2023), reiterating the specificity of ABI5 physical association with NUP1.

The protein level of ABI5 is known to be regulated by various post-translational modifications (Skubacz et al., 2016). Under abiotic stress, ABA activates ABI5 by phosphorylation (Nakashima et al., 2009; Wang et al., 2013). Under optimal conditions, ABI5 is ubiquitinated, which prepares it for degradation through the 26S proteasome pathway (Lopez-Molina et al., 2001; Stone et al., 2006; Liu and Stone, 2010). Our data suggest that ABI5 is degraded in nucleoplasm near NPC; however, two earlier studies proposed nuclear bodies (NB) to be the site of degradation. Lopez-Molina et al. (2003) suggested that ABI FIVE binding protein targets ABI5 for ubiquitin-mediated degradation in nuclear bodies. Similarly, Zhao et al. (2016) also proposed the same degradation site based on the CROWDED NUCLEI 3 colocalization with ABI5 in nuclear bodies. There are two possible reasons for this discrepancy. First, Lopez-Molina et al. (2003) used a *35S* promoter and Zhao et al. (2016) used a β-estradiol promoter to drive the expression of ABI5, which might not reflect the true expression and localization of the protein. We also tried using the *35S*-driven expression of *ABI5* for localization study and found it was distributed in all compartments of the nucleus (Supplementary Fig. S7), similar to that observed by Liu and Stone (2013). This ABI5 distribution pattern altered when we used the native promoter of *ABI5* for its expression (Fig. 7). Second, both studies did not track the timing of ABI5 expression and degradation. In this work, we optimized the timing of ABI5 expression and degradation with ABA treatment (Supplemental Fig. S6). Under our experimental conditions, ABI5 was induced after 4 hrs of ABA treatment (20 µM) and ABI5 could no longer be detected after another 6 hrs after removal of ABA, which indicates its degradation timeframe. Thus, observation of ABI5 at a different time may lead to different localization of ABI5. In a few cases, we also found some punctuate patterns of ABI5 indicating its localization in nuclear bodies (Supplemental Fig. S8), as reported by Lopez-Molina et al. (2003) and Zhao et al. (2016). However, we speculate that some kind of post-translational modification of ABI5 may result in its temporary retention in nuclear bodies/nucleolus. This is supported by Miura et al. (2009), which reported that ABI5 is sumoylated by SAP AND MIZ1 DOMAIN-CONTAINING LIGASE1 (SIZ1) and the sumoylated ABI5 is protected from ubiquitin-mediated degradation by promoting its localization in nuclear bodies. The punctuate pattern of ABI5 may be the condensate formed through phase separation. To evaluate the possibility of phase separation of ABI5, we used IUPred2 to determine the presence of any intrinsic disorder region (IDR), which is required for liquid-liquid phase separation. ABI5 is predicted to have an IDR between amino acids 355-405 (Supplemental Fig. S9), which may be vital for phase separation. Future work is needed to investigate whether phase separation indeed plays a role in the ABI5 accumulation in nucleolus/nuclear bodies.

Based on our data and the literature related to ABI5 degradation, we propose an ABI5 degradation model in *Arabidopsis* (Fig. 8). Under standard growth conditions, both wild type and *nup1-1* seeds show timely germination albeit the reduced efficiency of the proteasome-based nuclear degradation of ABI5 in *nup1-1*. This might be because of the less basal level of ABI5 in *nup1-1* compared to wild type (based on Fig 2A and 6E). However, during germination under stress conditions, *nup1-1* seeds have more ABI5 than Col-0 (based on Fig. 6F). Proteasomes in wild-type plants can efficiently degrade the increased level of ABI5 for timely germination. But, since NUP1-mediated proteasome activity is lowered in *nup1-1*, the rate of ABI5 degradation is slower. This leads to the accumulation of ABI5 and delayed germination. Increased ABI5 level can also self-activate its transcription through a positive feedback loop, which may help to explain the increased *ABI5* transcript level in *nup1-1* compared to Col-0 (Fig 4B). This self-regulation of ABI5 was previously reported (Xu et al., 2014). Thus, we propose, NUP1 indirectly reduces ABI5 transcription through directly promoting ABI5 protein degradation. The slow rate of ABI5 degradation in *nup1-1* under stress may lead to its retention in the nucleolus. This might be through post-translational modifications like sumoylation which was shown previously to promote its localization to nuclear bodies (Miura et al., 2009). To our knowledge, this is the first evidence in plants showing that NPC can maintain protein homeostasis through anchoring proteasome. Our work helps establish NPC as a site of protein degradation and surveillance. NPC is ideally positioned in the cell between the nucleus and cytoplasm to regulate protein level, mRNA export, and transcription. It would be interesting in the future to explore other proteins that are regulated by NPC tethered proteasome. In summary, our data strongly implicates NUP1 as a key modulator of seed germination through the ABA/ABI5 degradation pathway. Our results along with many others in this research area could be helpful in the future for breeding crops that can better germinate and grow under abiotic stress conditions, which is important, more than ever, for agriculture with the climate changes.

**Figure 8.**
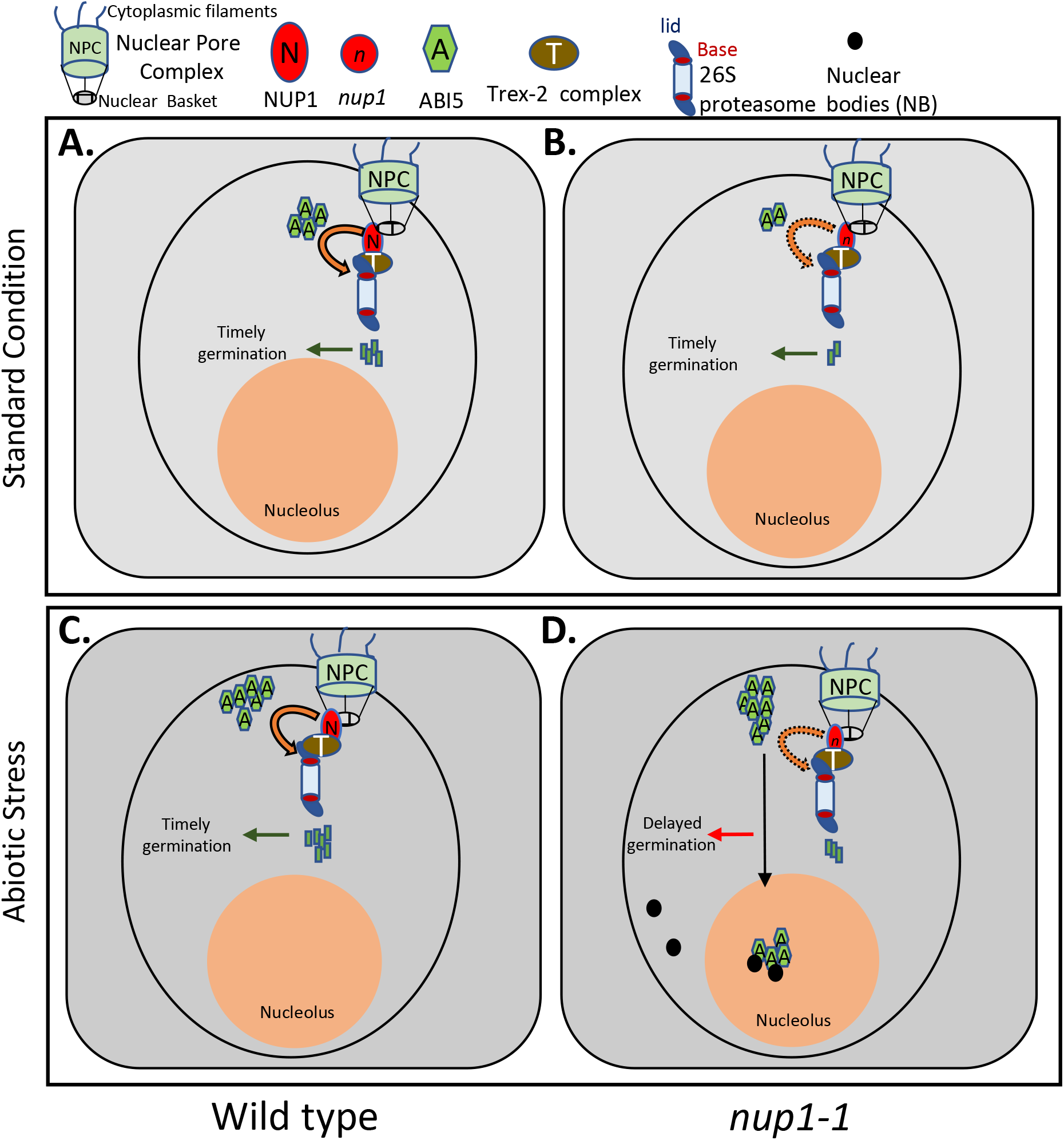
Proposed working model of proteasome-based ABI5 degradation near NPC during germination. A-B. Under standard growth condition (top panel). A. Col-0 wild-type seeds having a moderate amount of ABI5 degraded by the 26S proteasome-based degradation near NPC (indicated by curved arrow). B. *nup1-1* seeds have less amount of basal ABI5; and therefore, they also show timely germination despite the reduced activity of the proteasome (indicated by dotted curved arrow). C-D. Under abiotic stress such as ABA treatment (bottom panel). C. Col-0 seeds produce more ABI5, but it is degraded over time. D. *nup1-1* seeds accumulate more ABI5, but ABI5 degradation is slowed down due to reduced proteasomal activity, thus showing delayed germination (based on CHX assay Fig. 6G). Some ABI5 was seen as a bright spot in nucleolus indicating its localization in nuclear bodies (NB) (Supplementary Fig.8).

## Materials and Methods

### Plant materials and growth conditions

*Arabidopsis* wild type (Col-0) and the T-DNA mutants *nup1-1* (SALK_104728), *nup1-2* (SALK_020221), *nup1-3* (SAIL796H02), *abi1-2* (SALK_072009), *abi2-2* (SALK_015166C), *abi3-7* (SALK_038579), and *abi5-8* (SALK_013163) were all obtained from ABRC. The genotyping and qRT-PCR primers for characterization of these mutant lines are listed in Supplemental Table S7. Seeds were surface sterilized and placed in a dark room at 4°C for 2 d. Plants were grown in a walk-in growth room at 23°C, 150 μmol/m^2^/s light intensity, and 50 % humidity at 16 hrs light/8 hrs dark conditions. Double mutants were created by crossing single mutants and subsequent selection of double mutants by PCR-based genotyping. Leaves of three weeks-old tobacco (*Nicotiana benthamiana*) plants were used for transient expression assay.

### Germination test

For the germination assay, seeds were sterilized with 70 % ethanol for 10 min and washed with sterile water. Then seeds were sown on ½ Murashige and Skoog (MS) medium plates (1.5 % sucrose, 0.8 % agar, pH 5.75) with or without various concentrations of supplements; ABA (for abiotic stress), NaCl (for salt stress), and sorbitol (for osmotic stress). The seeds were vernalized in a dark room at 4°C for two days and then transferred to the growth room. The ratio of radicle emergence and cotyledon emergence was calculated every day. Cotyledon emergence is considered successful germination in this study. At least 50 seeds per biological replicate and three independent replicates were conducted. Radicle emergence or cotyledon emergence was calculated on the 5th day of imbition to determine any statistical differences.

### Drought assay

For the drought assay, 10-day-old Col-0 and *nup1-1* seedlings grown on ½ MS medium plates were left open for drying in a laminar flow hood for six hrs. Then seedlings were put back in the regular growth room for the next 5 days. Then, the percentage of survived plants was calculated.

### Plasmid construction and transgenic plants

The transgenic line expressing *NUP1* fused with *YFP* driven by its native promoter in the *nup1-2* background (*nup1-2 pNUP1::NUP1-YFP*) was previously described (Lu et al., 2010). ABI5 genomic DNA with or without its native promoter (∼ 2 kb) was amplified from a bacterial artificial chromosome (BAC) clone (BAC F2H17). The amplified DNA fragments were cloned into the D-TOPO vector (Cat no #K240020, ThermoFisher Scientific) and then inserted into the PMDC107 and PMDC84 vectors through LR cloning (Gateway technology). The created plasmid DNAs were later sequenced for accuracy. After sequencing, the plasmid constructs were introduced to *Agrobacterium tumefaciens* (GV3101) through electroporation. *Agrobacterium*-mediated transformation of *Arabidopsis* in the *abi5-8* and *nup1-1 abi5-8* backgrounds was performed using the floral dip method (Clough and Bent, 1998). Transgenic lines expressing *ABI5* with the native promoter in different genetic backgrounds, (*abi5-8 pABI5::ABI5-GFP*) and (*nup1-1 abi5-8 pABI5::ABI5-GFP*), were used for ABA related studies. The plasmid expressing *35S::ABI5-CFP* was used for the co-localization assay. The *NUP1* overexpression *Arabidopsis* line *(nup1-2 35S::NUP1-YFP*) is characterized in Supplemental Fig. S3.

### Protein extraction and Western blot

Protein extraction and Western blot were performed as described with slight modifications (Shu et al., 2021). Seeds and seedlings (0.5-2 gm) were ground in liquid nitrogen and proteins were extracted with lysis buffer containing 50 mM Tris-HCl, 10 mM ethylenediaminetetraacetic acid (EDTA), 1× complete™ Protease Inhibitor Cocktail (Roche), 0.1 mM phenylmethylsulfonyl fluoride (PMSF), and 1 % sodium dodecyl sulfate (SDS) (w/v). The protein solution was centrifuged at 4°C for 15 min to remove cellular debris. Then, the protein preparation was dissolved in a 3x SDS loading buffer and 20-30 µl of the protein preparation was loaded onto an SDS polyacrylamide gel (10%). The protein size was separated in the gel for 2 hrs at 120 V and transferred to a polyvinylidene fluoride (PVDF) membrane. Then, 5 % (w/v) non-fat milk/Tris-buffered saline (TBS) was used to block the membrane for 1hr by gentle shaking at room temperature. After that, the primary antibody (anti GFP: Abcam, ab290, 1:20,000 dilution; anti-ABI5 antibody: Abcam (Ab98831) at 1:1000 dilution) was added and incubated overnight at 4°C. The next day, the membrane was washed with TBS and a Horse Radish Peroxidase conjugated secondary antibody (A0545, Sigma) was added and incubated for an hour at room temperature. The excess secondary antibody was removed by washing three times with TBS buffer. The protein bands were visualized by using the Western Blotting Reagents (RPN2106, GE Healthcare) and chemiluminescence was captured using MicroChemi (DNR Bio-Imaging System).

### RNA isolation

RNAs were isolated from seeds and seedlings. Seeds (50 mg) were ground in liquid nitrogen and extraction buffer (0.4 M LiCl, 0.2 M Tris pH 8, 25 mM EDTA, and 1 % SDS), and chloroform was added. The solution was centrifuged for 3 min at 4°C at 10,000 RPM and the supernatant was transferred to a new Eppendorf tube and 500 µl of water-saturated acidic phenol was added and vortexed thoroughly. The solution was then centrifuged for 3 min at 4°C. The supernatant was transferred to a new tube and 1/3 volume of 8 M LiCl was added. The supernatant was then precipitated at −20°C for 3 hr and spun for 30 min at 4°C to form a pellet. 470 µl of water was added to dissolve the pellet. RNA was again precipitated with 3M sodium acetate, and absolute ethanol and centrifuged for 10 min to remove carbohydrates. The supernatant was again precipitated in ethanol, dissolved in water, and treated with DNAse. RNA from young seedlings was isolated using the protocol from Shu et al. (2019) using the RNAeasy Plant Mini Kit (Qiagen) following the manufacturer’s protocol.

### RNA-Seq and data analysis

RNA sequencing and analysis were done as described in Chen et al. (2019). Briefly, total RNA was isolated, purified, and sent for deep sequencing at Novogene, China. Each sample had three biological replicates. The obtained reads were cleaned by removing low-quality sequences by using the Trimmomatic program. Those reads were mapped to the TAIR10 *Arabidopsis* genome using TopHat v2.0.4 with default setting except that a minimum intron length of 20 bp and a maximum intron length of 4,000 bp were used. DEGs were calculated using the Cufflinks package (Trapnell et al., 2012). In short, the alignment files after running TopHat were used as input to Cufflink to generate a transcriptome assembly for each sample. All these assemblies were merged using the Cuffmerge. The reads and merged assembly were used as input in Cuffdiff for the calculations of DEGs. Genes with at least a 2-fold change in expression (FDR ≤ 5 %, p-value < 0.05) were considered to be differentially expressed. The gene ontology (GO) enrichment analysis was performed by online tools Agri-go (http://bioinfo.cau.edu.cn/agriGO/). Heat maps and volcano plots were generated using the R software.

### Gene Expression Analysis

Total RNA was extracted from *Arabidopsis* seeds and seedlings of Col-0 and mutant plants as described above. For estimating gene expression, 500 ng of total RNA was used for reverse transcription using the SuperScript III Reverse Transcriptase kit (Invitrogen). Real-time PCR was performed using the iQ SYBR Green supermix (Bio-Rad) on a CFX96 real-time PCR machine (Bio-Rad). Gene expression was normalized to *ACTIN 2* (*ACT2*). All primers used are listed in Supplemental Table S7.

### BiFC Assay

Full-length *NUP1* and *ABI5* coding sequences of *Arabidopsis* were amplified and recombined into the pDONR221 vector by BP reaction (Gateway technology). Those vectors with insert were sequenced to ensure no errors were introduced during PCR amplification. The entry vector pDONR221 with insert was then transferred into the modified pEarleyGate 201-N-YFP or pEarleyGate 202-C-YFP vector by LR reaction (Gateway technology) (Lu et al., 2010). The final constructs were introduced into *Agrobacterium tumefaciens* strain GV3101 and the resulting bacteria were used to infiltrate the abaxial side of tobacco (*Nicotiana benthamiana*) leaves. After 2-3 days, the fluorescence signals were detected for positive protein interactions, and no signals were detected when two proteins did not interact. pEarleyGate 201-N*-NUP1-YFP* and pEarleyGate 202-C-CENTRIN2 were used as negative controls. The imaging of the fluorescence was done using a confocal microscope (Olympus FV1000). Sequences of the primers used are listed in supplemental Table S7.

### Nuclear staining and Confocal microscopy

For fluorescence analysis, *Arabidopsis* seedlings were stained with 10 µg/ml propidium iodide (PI) and/or 5 µg/ml 4′, 6-diamidino-2-phenylindole (DAPI) (Sigma #D9542) for the indicated times in dark at room temperature. The seedlings were washed with distilled water (ddH_2_O) for 15 min to remove excess chemicals. The treated seedlings were immediately imaged with a confocal microscope (Olympus FV 1000). The following conditions were used for the detection of various signals: for GFP detection, excitation 488 nm, emission 510 nm; for YFP detection, excitation 515 nm, emission 520-550 nm; for DAPI, excitation 405 nm, emission 470-500 nm; for CFP detection, excitation 405 nm; emission 485 nm; for PI detection, excitation 535 nm; emission 615 nm.

### Co-localization assay

To detect the co-localization of ABI5 and NUP1, *Agrobacterium* expressing *35S::ABI5-GFP* was transiently expressed in the stable *Arabidopsis* transgenic line *nup1-2 pNUP1::NUP1-YFP*. After two days, the transiently transformed leaves were analyzed by confocal microscopy. Cells expressing both GFP and YFP proteins were examined for the co-localization pattern of two proteins.

### Statistical Analysis

Means, standard deviations (SD), and standard error of the mean (SEM) were calculated using Excel 2016 (Microsoft Corp., Redmond, Washington). All experiments were performed using at least three biological replicates and additional three technical replicates were included in gene expression analysis by qRT-PCR. SD was used for the analysis of experiments with biological replicates and SEM was used for the analysis of experiments with biological and technical replicates. Statistical analysis was performed by Student’s test or ANOVA. A p-value of 0.05 or less was considered as a statistically significant difference indicated by * and a p-value of 0.01 or less was denoted by **.

### Accession Numbers

The TAIR accession numbers for the sequences of genes mentioned in this study are as follows: *NUP1* (At3g10650), *ABI1* (AT4G26080), *ABI2* (AT5G57050), *ABI3* (AT3G24650), *ABI4* (AT2G40220), and *ABI5* (AT2G36270).

## ACKNOWLEDGMENTS

We thank the *Arabidopsis* Biological Resource Centre (ABRC) for providing the mutant seeds used in this study. This work was supported by grants from the Natural Science and Engineering Research Council of Canada (RGPIN/04625-2017, to Y.C.); and Agriculture and Agri-Food Canada (to Y.C.).

## Author Contributions

YC conceived the project. YC, RKT, GT, and SEK designed the experiments. GT contributed to BiFC assay. RKT, JShu, CC, and CL analyzed the RNA-Seq data. XX, JSong, YY and SB did the crossing and genotyping. VN performed all the Sanger DNA sequencing. JL performed a qRT-PCR experiment. RKT and ML conducted germination assay. RKT performed all the rest of the experiments. RKT and YC wrote the manuscript. All authors read and approved the final article.

## DATA AVAILABILITY STATEMENT

RNA-Seq data have been deposited in the National Center for Biotechnology Information GEO database under accession number GSE189513. Materials are available from the corresponding author upon request.

