## Supplementary figures for "NUCLEOPORIN1 mediates proteasome-based degradation of ABI5 to regulate *Arabidopsis* seed germination"

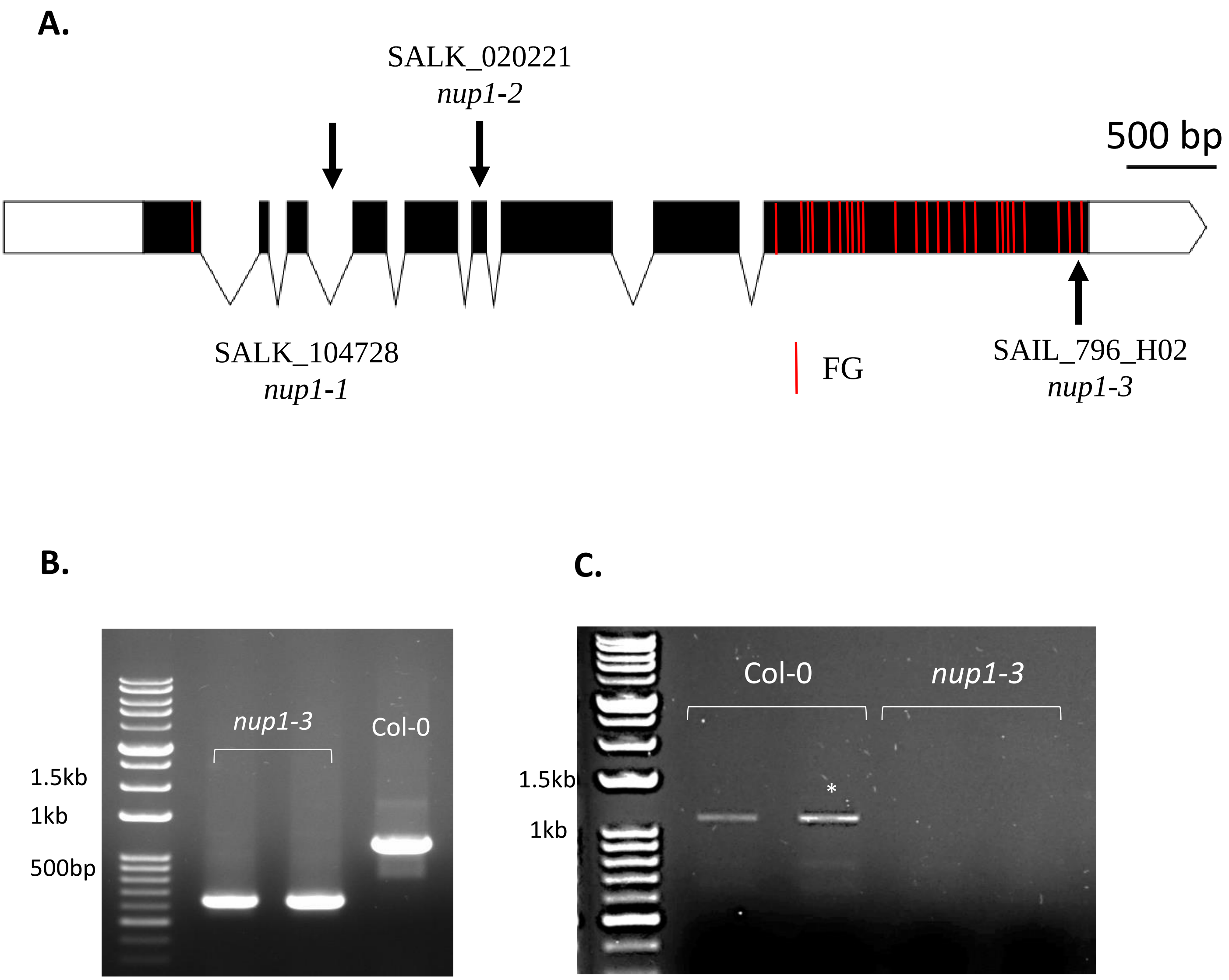
A.
SALK_020221
nup1-2
500 bp
SALK_104728
nup1-1
FG
SAIL_796_H02
nup1-3
B.
nup1-3
Col-0
1.5kb
1kb
500bp
C.
Col-0
nup1-3
1.5kb
*
1kb

**Supplementary Figure 1. Characterization of NUP1 T-DNA insertion lines**

A. Structure of the *NUP1* gene showing T-DNA insertion sites. The *NUP1* gene structure shows the introns (V-shaped lines), exons (black boxes), and untranslated regions (empty boxes). The red vertical lines indicate the FG repeats. The black arrow indicates the sites of T-DNA insertion. The *nup1-1* (SALK_104728), *nup1-2* (SALK_020221) *nup1-3* (SAIL_796_H02) have a T-DNA insertion in the third intron, sixth exon and ninth exon, respectively. Scale bar: 500 bp.

B. Genotyping of *nup1-3* allele shows single band around 700 bp indicating homozygous allele compared to Col-0 wild-type which have single band around 1kb.

C. Expression of NUP1 in *nup1-3* allele. qPCR and subsequent product ran in agarose gel shows no product in *nup1-3* allele.

**A.**

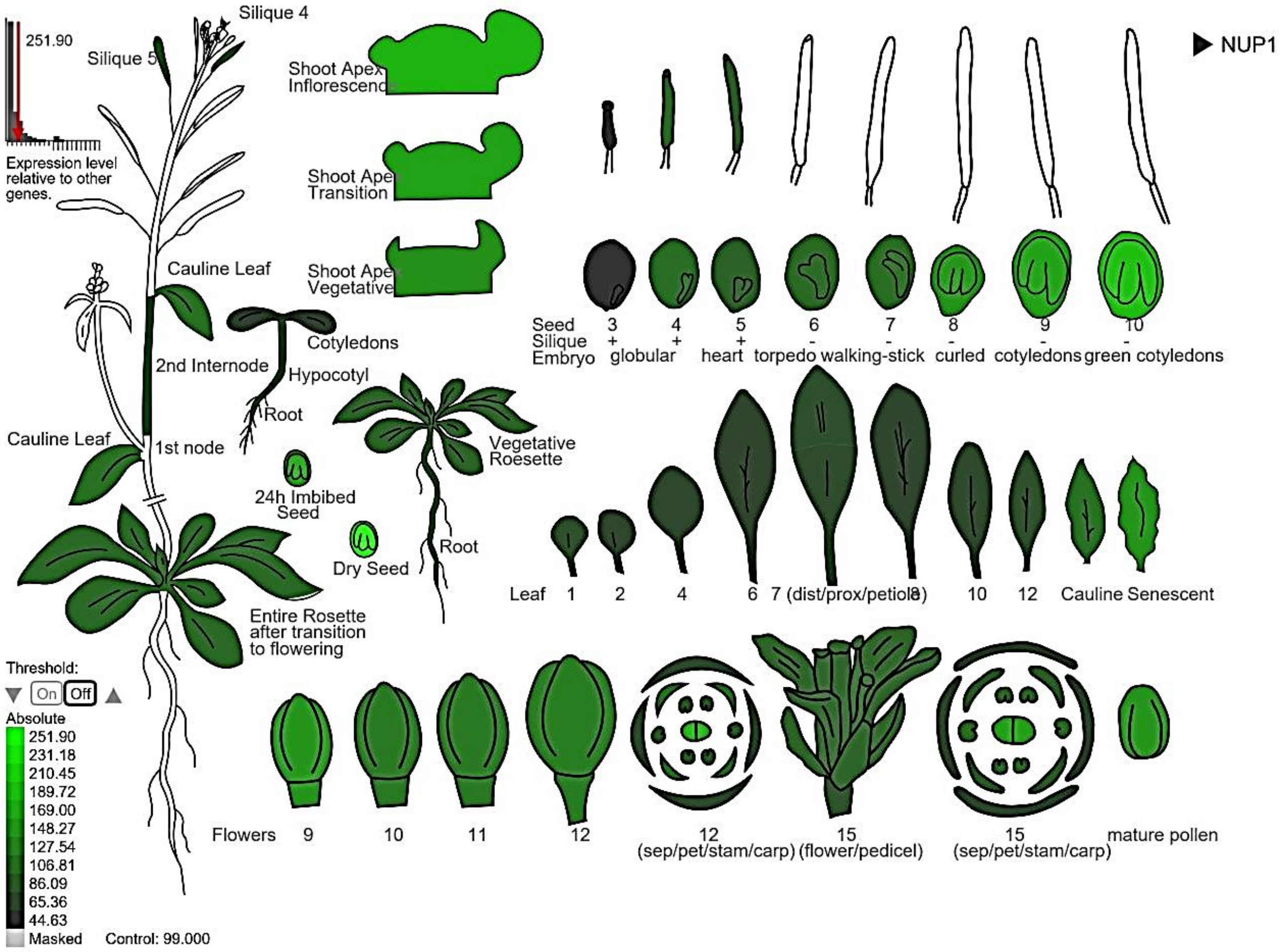

NUP1
251.90
Expression level relative to other genes.
Silique 4
Silique 5
Shoot Apex Inflorescence
Shoot Apex Transition
Shoot Apex Vegetative
Cauline Leaf
2nd Internode
Cotyledons
Hypocotyl
Root
Cauline Leaf
1st node
24h Imbibed Seed
Dry Seed
Vegetative Roesette
Root
Entire Rosette after transition to flowering
Seed 3 4 5 6 7 8 9 10
Silique + + + - - - - -
Embryo globular heart torpedo walking-stick curled cotyledons green cotyledons
Leaf 1 2 4 6 7 (dist/prox/petiole) 8 10 12 Cauline Senescent
Threshold:
On Off
Absolute
251.90
231.18
210.45
189.72
169.00
148.27
127.54
106.81
86.09
65.36
44.63
Masked
Control: 99.000
Flowers 9 10 11 12
12 (sep/pet/stam/carp)
15 (flower/pedicel)
15 (sep/pet/stam/carp)
mature pollen


**B.**

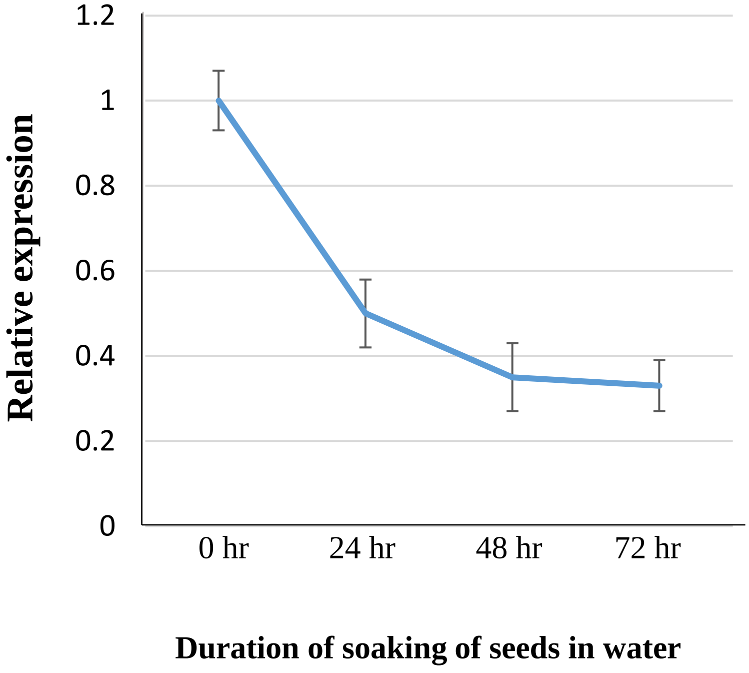

Relative expression
1.2
1
0.8
0.6
0.4
0.2
0
0 hr
24 hr
48 hr
72 hr
Duration of soaking of seeds in water

**Supplementary Figure 2. *NUP1* gene expression pattern in *Arabidopsis thaliana*.**

**A.** *NUP1* gene expression in different tissues under standard growth conditions of *Arabidopsis* across various development stages. Data were downloaded from the efp browser http://bar.utoronto.ca/efp2/*Arabidopsis*/*Arabidopsis*_eFPBrowser2.html (Winter et al., 2007). Gene expression levels are shown in the color key; dark green is the lowest, light green is the highest, and white represents no gene expression. **B.** Expression of *NUP1* in dry and imbibed seeds. Dry seeds of Col-0 were soaked in water for the indicated times and *NUP1* gene expression was measured by qRT-PCR. The gene expression was normalized to *ACTIN2*. Values are mean ± SEM from three biological and three technical replicates. The value for 0 hr was set to 1 for comparison.

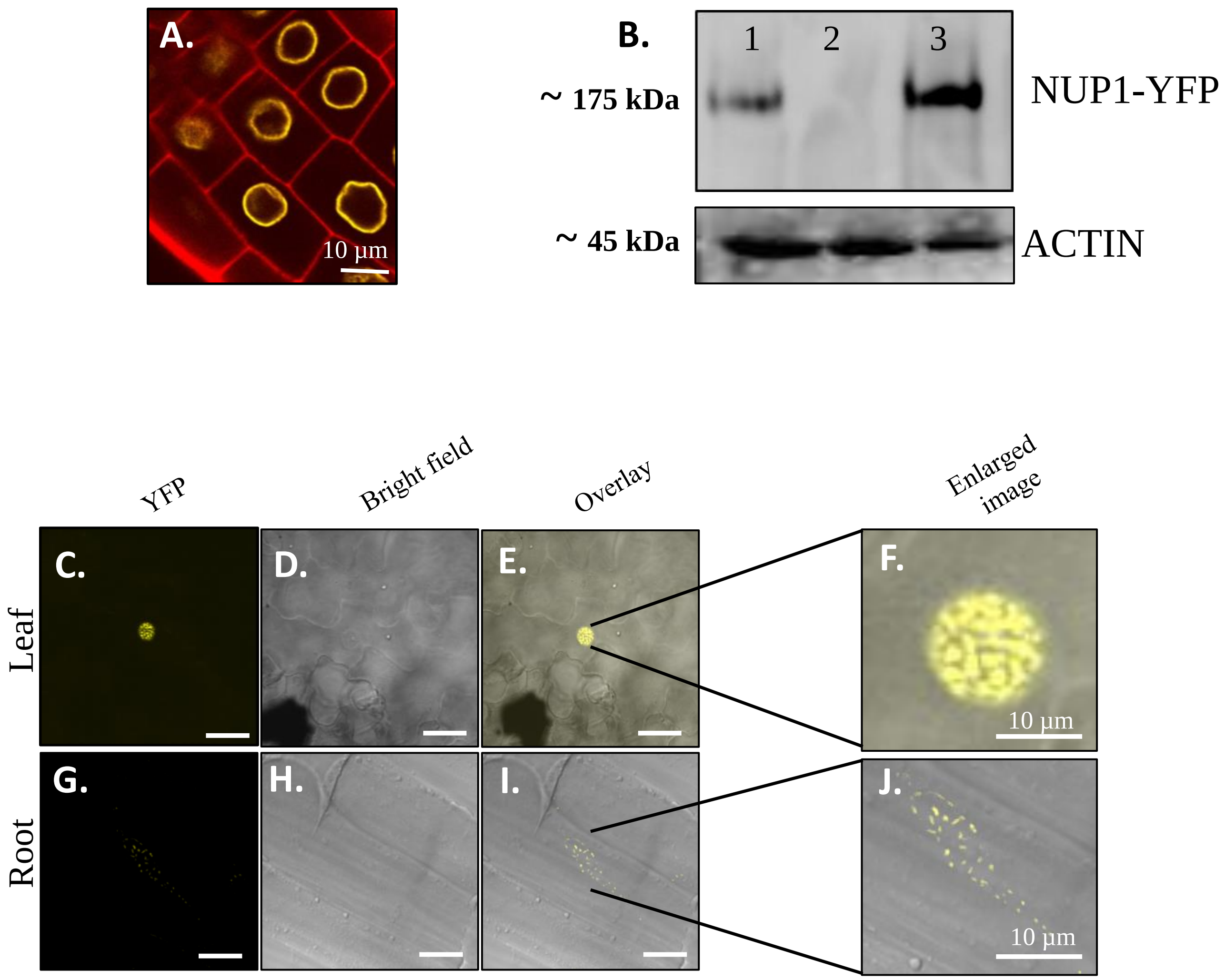
A.
10 µm
B.
1
2
3
~ 175 kDa
NUP1-YFP
~ 45 kDa
ACTIN
YFP
Bright field
Overlay
Enlarged image
Leaf
C.
D.
E.
F.
10 µm
Root
G.
H.
I.
J.
10 µm

**Supplementary Figure 3. Characterization of an *Arabidopsis* transgenic line overexpressing *NUP1* driven by the *35S* promoter.**

**A.** Normal localization of wild-type level NUP1 (used as a control) in root cells of 7-day-old transgenic seedlings expressing NUP1-YFP under its native promoter (*nup1-2 pNUP1::NUP1-YFP*). NUP1 is localized around nuclei in nuclear envelope regions forming a circular shape. PI (red) was used to stain the cell wall. Scale bar: 10 µm. **B.** Western blot analysis of proteins isolated from the transgenic line overexpressing *NUP1*. Lane 1: The transgenic line *nup1-2 pNUP1::NUP1-YFP* as positive control; lane 2: Col-0 as a negative control; lane 3: The NUP1-overexpressing transgenic line (*nup1-2 35S::NUP1-YFP*). The anti-GFP antibody (Abcam, ab290) was used to detect NUP1 and the anti-ACTIN antibody (ab197345) was used to detect ACTIN (loading control). **C-F.** NUP1 accumulation in the nucleus of NUP1-overexpressed transgenic line's leaves. **C:** YFP signal; **D:** bright-field image; **E:** overlay of C and D; and **F:** an enlarged section of E showing nuclear accumulation of NUP1. Scale bar: 10 µm. **G-J.** NUP1 accumulation in the nucleus of NUP1-overexpressing transgenic roots. **G:** YFP signal; **H:** bright field image; **I:** overlay; and **J:** an enlarged section of **I** showing nuclear accumulation of NUP1. Scale bar: 10 µm.

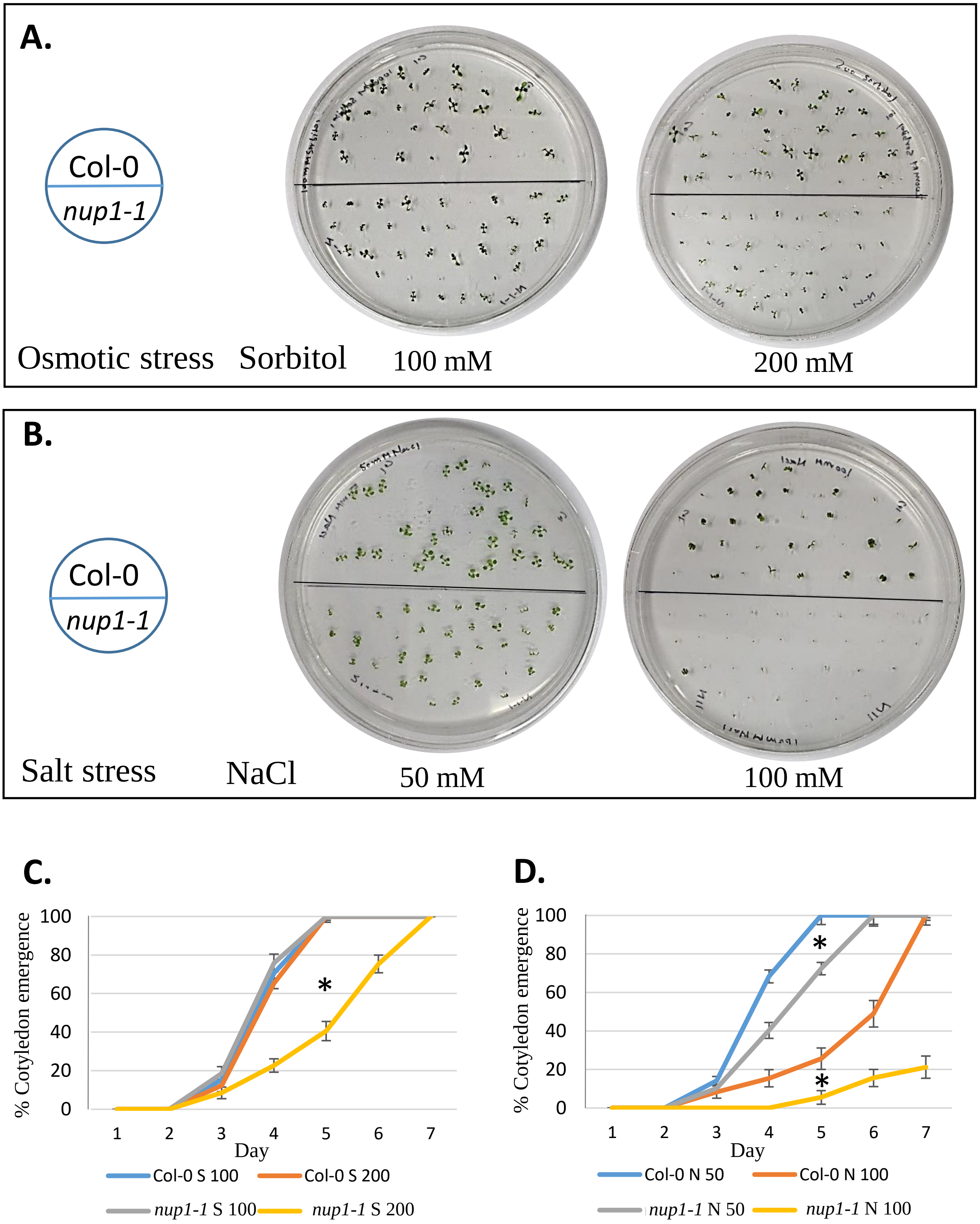
A.
Col-0
nup1-1
Osmotic stress
Sorbitol
100 mM
200 mM
B.
Col-0
nup1-1
Salt stress
NaCl
50 mM
100 mM
C.
% Cotyledon emergence
100
80
60
40
20
0
1
2
3
4
5
6
7
Day
*
Col-0 S 100
Col-0 S 200
nup1-1 S 100
nup1-1 S 200
D.
% Cotyledon emergence
100
80
60
40
20
0
1
2
3
4
5
6
7
Day
*
*
Col-0 N 50
Col-0 N 100
nup1-1 N 50
nup1-1 N 100

**Supplementary Figure 4. The response of *Arabidopsis nup1-1* seed germination to exogenous sorbitol and salt stress.**

**A.** Seeds germinated on ½ MS medium plates with sorbitol (100 and 200 mM) to mimic osmotic stress. **B.** Seeds germinated on ½ MS medium plates with NaCl (50 and 100 mM) to mimic salt stress. **C-D.** Percentage of cotyledon emergence under osmotic stress (C**),** and salt stress (D**).** Sorbitol = S and NaCl = N in the graphs C and D. The number followed by S and N is the concentration in millimolar (mM). The values are mean ± SD. Statistically significant differences were determined by Student's t-test from day 5 data, $*p < 0.05$.

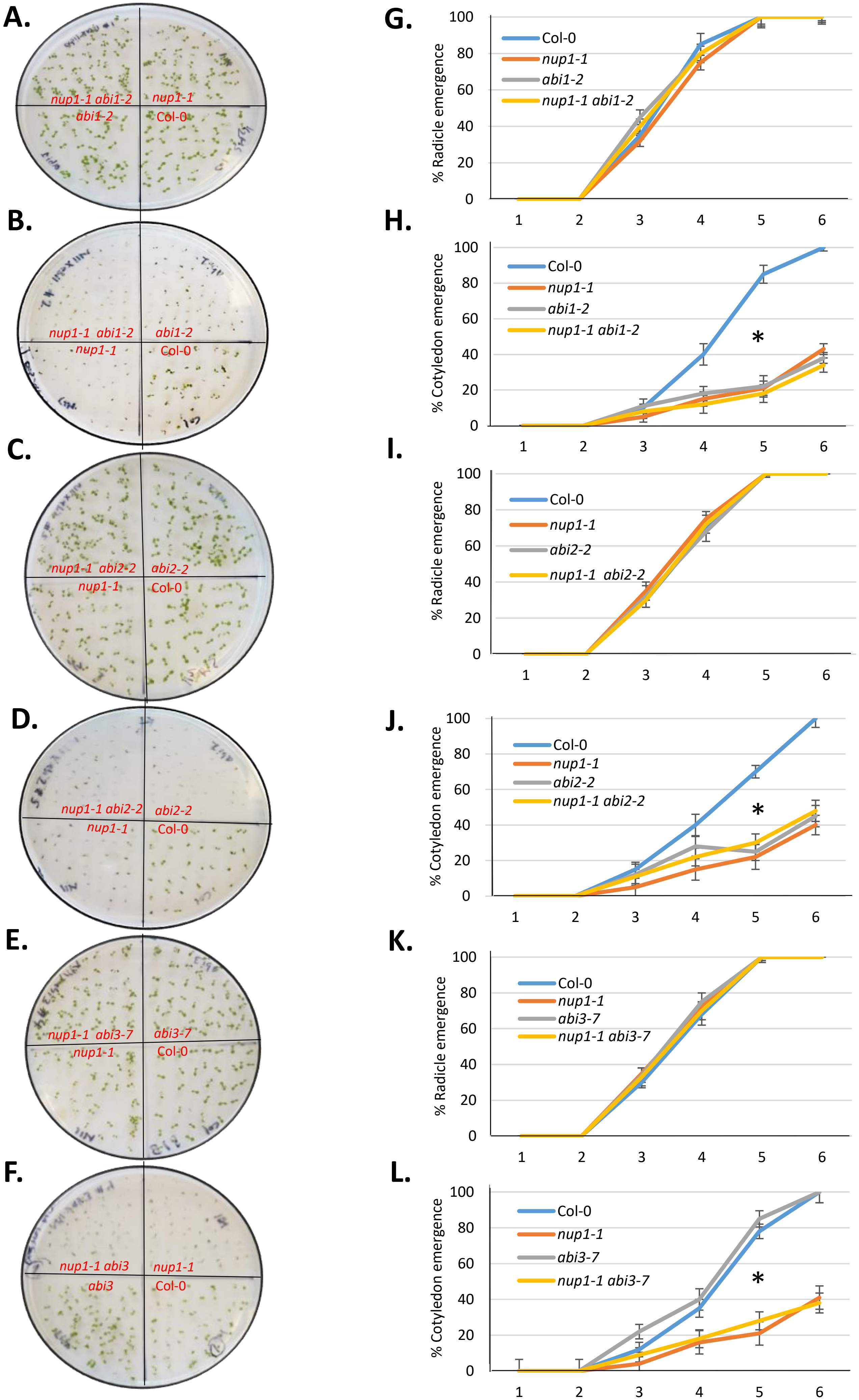
A.
nup1-1 abi1-2
nup1-1
abi1-2
Col-0
B.
nup1-1 abi1-2
abi1-2
nup1-1
Col-0
C.
nup1-1 abi2-2
abi2-2
nup1-1
Col-0
D.
nup1-1 abi2-2
abi2-2
nup1-1
Col-0
E.
nup1-1 abi3-7
abi3-7
nup1-1
Col-0
F.
nup1-1 abi3
nup1-1
abi3
Col-0
G.
% Radicle emergence
Col-0
nup1-1
abi1-2
nup1-1 abi1-2
H.
% Cotyledon emergence
Col-0
nup1-1
abi1-2
nup1-1 abi1-2
*
I.
% Radicle emergence
Col-0
nup1-1
abi2-2
nup1-1 abi2-2
J.
% Cotyledon emergence
Col-0
nup1-1
abi2-2
nup1-1 abi2-2
*
K.
% Radicle emergence
Col-0
nup1-1
abi3-7
nup1-1 abi3-7
L.
% Cotyledon emergence
Col-0
nup1-1
abi3-7
nup1-1 abi3-7
*
0 20 40 60 80 100
1 2 3 4 5 6

**Supplementary Figure 5. Mutations in *ABI1*, *ABI2*, or *ABI3* cannot rescue the ABA-sensitive germination phenotype of *nup1-1*.**

**A-F.** Seed germination, recorded at day 5, of the genotypes indicated with (B, D, F) or without 1 µM ABA treatment (A, C, E). **G-L.** Graph showing the percentage of radicle emergence (G, I, K) and cotyledon emergence (H, J, L) of seeds from different genetic backgrounds grown on ½ MS medium plates with 1 µM ABA treatment. The seed germination response without ABA treatment (A, C, E) served as control. There was no difference in radical emergence ratio among genotypes (G, I, K); however, the cotyledon emergence (germination) was delayed in *nup1-1*, *abi1-1*, *abi2-2*, and all the double mutants (*nup1-1 abi1-1*, *nup1-1 abi2-2*, and *nup1-1 abi3-7*) compared to Col-0.

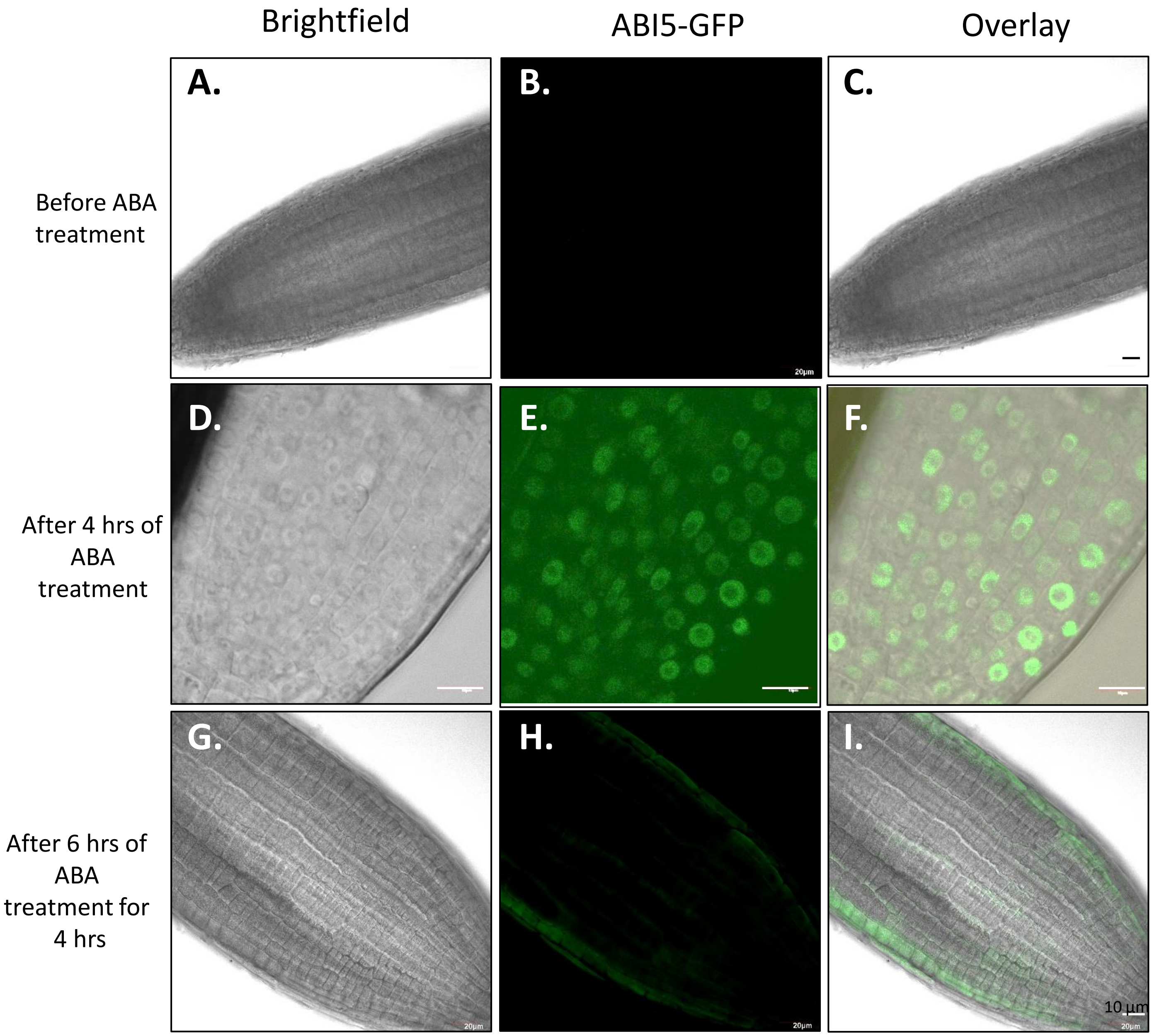
Brightfield
ABI5-GFP
Overlay
A.
B.
C.
D.
E.
F.
G.
H.
I.
Before ABA treatment
After 4 hrs of ABA treatment
After 6 hrs of ABA treatment for 4 hrs
20µm
10 µm

**Supplementary Figure 6. Tracking the localization of ABI5 in *Arabidopsis* roots.**

*Arabidopsis* transgenic line expressing ABI5-GFP (*abi5-8 pABI5::ABI5-GFP*) was treated with ABA (20 µM) for 4 hrs. Then, subcellular localization of ABI5 was tracked at the indicated time A, D, and G: bright field: B, E, and H: GFP channel; and C, F, and I: merged images. Scale bar: 10 µm. **A-C.** The lateral section of a 5-day-old root was immediately observed before ABA treatment. **D-F.** The lateral section of a 5-day-old root was observed after 4 hrs of ABA treatment. **G-I.** The lateral section of a 5-day-old root was observed 6 hrs after the 4 hrs of ABA treatment.

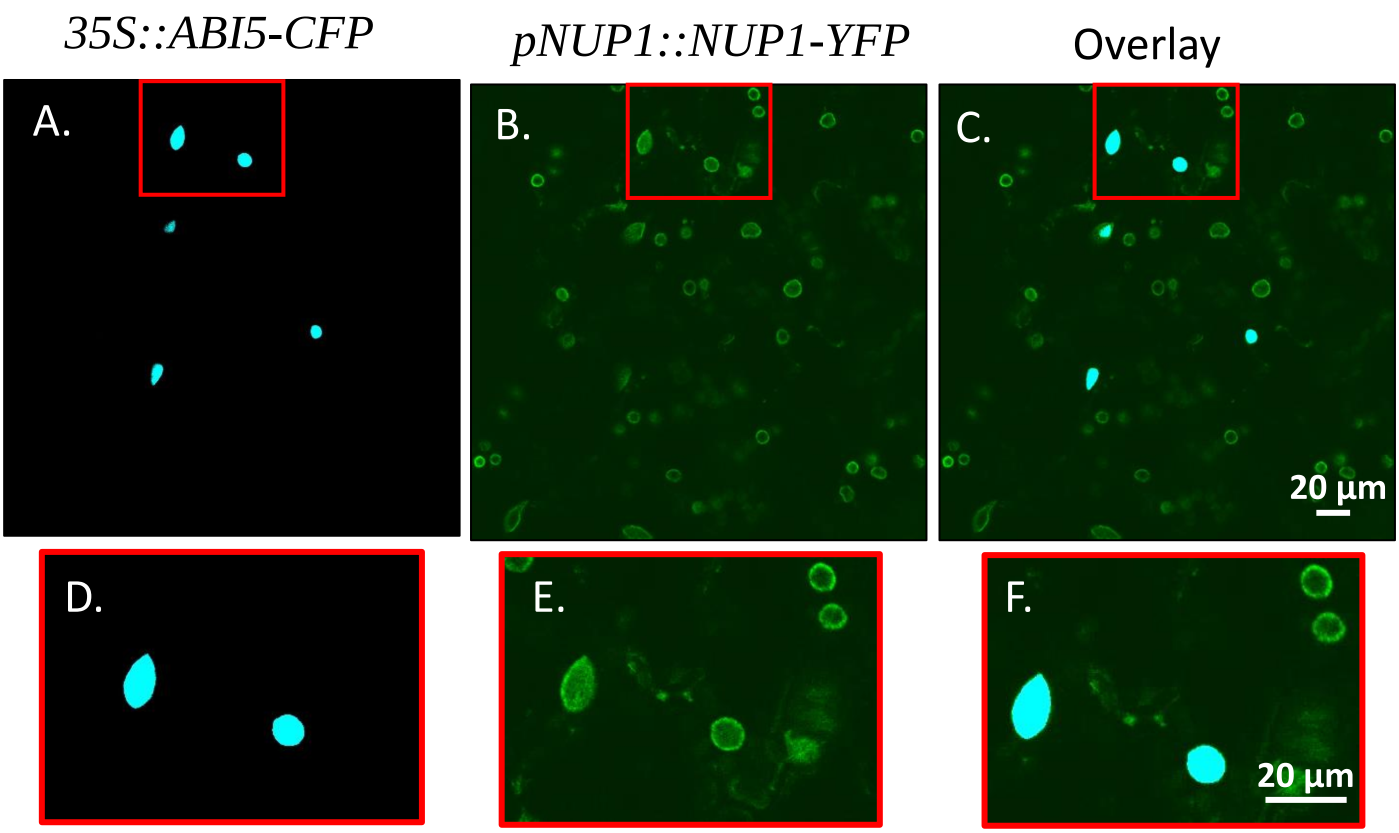

35S::ABI5-CFP
pNUP1::NUP1-YFP
Overlay
A.
B.
C.
20 µm
D.
E.
F.
20 µm

**Supplementary Figure 7. Subcellular localization of ABI5 in an *Arabidopsis* transgenic line over-expressing ABI5.**

The *Arabidopsis* transgenic line expressing *pNUP1::NUP1-YFP* was transformed with the plasmid over-expressing ABI5-CFP (*35S::ABI5-CFP*). **A.** ABI5-CFP; **B.** NUP1-YFP; and **C.** Merged. **D-F** is the enlarged version of the sections highlighted in red boxes in A-C.

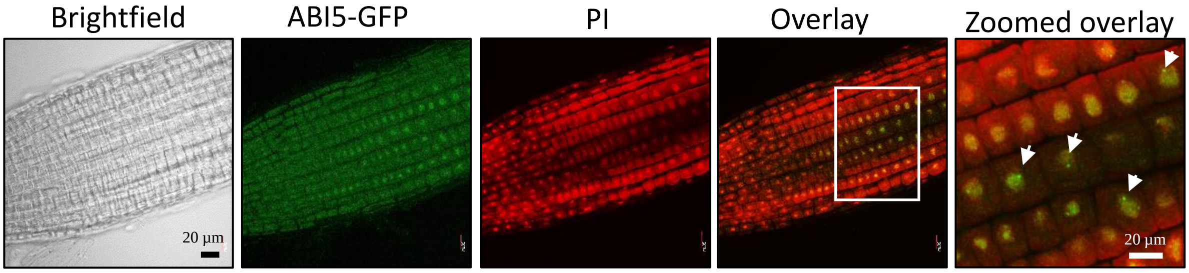

Brightfield
ABI5-GFP
PI
Overlay
Zoomed overlay
20 μm
20 μm

**Supplementary Figure 8. Expression of ABI5 under its native promoter in *Arabidopsis* roots showing its accumulation in nuclear bodies in the nucleolus.**

PI was used to label the nucleolus and/or cell walls. Bright green spots (condensate) shown by the white arrows are ABI5 proteins in nuclear bodies. Scale bar: 20 μm.

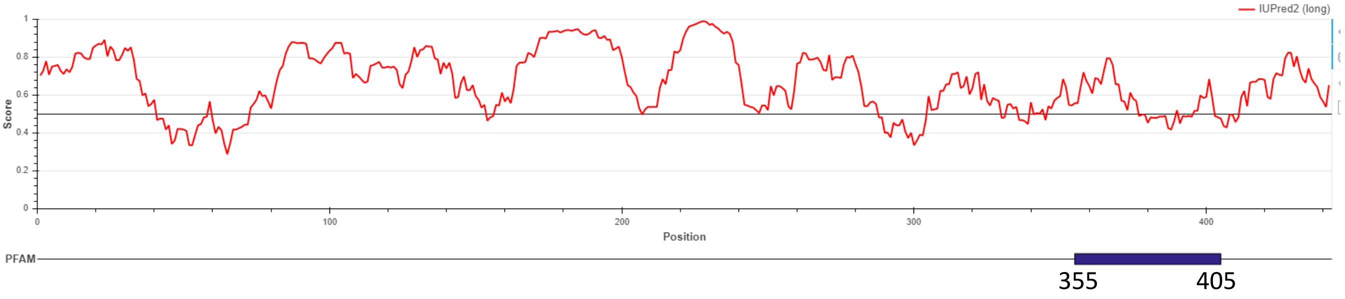

IUPred2 (long)
Score
Position
PFAM
355
405

**Supplementary Figure 9. Computational prediction of intrinsic disorder region of *Arabidopsis* ABI5 using IUPred2.**

A disorder region of ABI5 was predicted between amino acids 355-405, using the IUPred2 (long) option. This disorder region may be required for ABI5 condensate formation through phase separation. IUPred2 can be accessed at: https://iupred2a.elte.hu/plot_new
